# BioGraphX: Bridging the Sequence-Structure Gap via Physicochemical Graph Encoding for Interpretable Subcellular Localization Prediction

**DOI:** 10.64898/2026.01.21.700873

**Authors:** Abubakar Saeed, Waseem Abbas

## Abstract

Computational approaches for protein subcellular localization prediction are important for understanding cellular mechanisms and developing treatments for complex diseases. However, a critical limitation of current methods is their lack of interpretability: while they can predict where a protein localizes, they fail to explain why the protein is assigned to a specific location. Moreover, understanding protein behavior traditionally requires knowledge of three-dimensional structure, which is a costly and time-consuming process. Here, we propose BioGraphX, a novel encoding framework that constructs protein interaction graphs directly from protein sequences using biochemical rules. This approach provides a constraint-based structural proxy directly from sequence, reducing the dependency on experimentally determined three-dimensional structures. Building upon this representation, BioGraphX-Net demonstrates superior performance on the DeepLoc 2.0 benchmark by integrating ESM-2 embeddings with the proposed features via a gating mechanism. Gating analysis shows that although ESM-2 embeddings provide strong contributions, BioGraphX features function as high-precision filters. SHAP analysis reveals feature importance patterns consistent with a sophisticated biophysical logic: sequence signals act as universal exclusion filters, while organelle-specific combinations of biophysical features enable precise compartment discrimination. Notably, Frustration features help resolve targeting ambiguities in complex compartments, reflecting evolutionary constraints while preventing mislocalization from sequence mimicry. It has the additional advantage of promoting Green AI in bioinformatics, achieving performance comparable to the state-of-the-art while maintaining a minimal parameter count of 13.46 million. In summary, BioGraphX not only provides accurate predictions but also offers new insights into the language of life.

## Introduction

Proteins are the most important part of our understanding of living things. Figuring out where they are in the cell is a key part of cellular processes, and it is very important for making sure precise drug discovery and comprehension of disease pathogenesis [1, 2]. In recent years, high-throughput sequencing technologies have advanced rapidly, generating enormous volumes of genetic data. This has led to an explosive increase in the number of discovered protein sequences. The immediate outcome of this imbalance is a vast and growing dark matter of uncharacterized proteins, where the rate of sequence discovery far outpaces our capacity for manual annotation.

Historically, identifying a protein’s subcellular destination has required direct experimental observation using techniques like fluorescence microscopy and immunoelectron microscopy [3]. These methods remain the gold standard for real-time, high-quality protein distribution analysis but are limited by high cost and specialized equipment requirements. In the post-genomic era, a widening gap between reviewed and unreviewed entries in the UniProt database, of which only a small fraction (less than 0.3%) has been manually curated in Swiss-Prot, has driven the development of computational methods to bridge this gap [4].

Over time, computational approaches to protein localization have evolved from simple motif-based methods to more advanced deep learning models. Initial studies in this field relied on conserved targeting signals, before gradually incorporating hand-crafted physicochemical features [5]. Several methods have been introduced to enhance localization predictions by integrating different types of biological data. YLoc+, for instance, predicts multiple locations by combining sorting signals [6], PROSITE [7] patterns, and Gene Ontology (GO) terms, whereas Fuel-mLoc [8] uses only GO terms from the ProSeq-GO database to identify protein locations in diverse organisms.

A major milestone was achieved with DeepLoc 1.0 [9], which was among the first methods to apply recurrent neural networks (RNNs) to protein sequence modeling. More recently, there has been growing interest in protein language models (pLMs), such as ESM-2 [10] and ProtT5 [11]. Although most protein language models depend on extensive self-supervision, LAProtT5 [12] introduced a sequence-focused deep learning approach for localization prediction. This transition led to the development of DeepLoc 2.0 [2], which delivered significant gains in global accuracy. More recently, LocPro [13] has emerged as a promising method that overcomes several limitations of sequence-only embeddings. LocPro introduces a dual-channel architecture that integrates global protein representations from the ESM-2 language model with 1,484 expert-driven descriptors from PROFEAT [14]. By restructuring these expert features into similarity-based images (PF-IMG) and processing them through a hybrid CNN-BiLSTM network, LocPro has achieved new state-of-the-art results in multi-label prediction across 91 subcellular localization labels spanning varying levels of granularity.

However, despite the success of sequence-driven hybrids like LocPro, there remains a critical gap in explicitly modeling the network of intramolecular interactions, such as hydrophobic packing, salt bridges, and hydrogen bonding, that govern protein folding. While LocPro relies on statistical sequence descriptors, BioGraphX-Net shifts the paradigm toward structural graph encoding. By representing proteins as interaction networks, our framework captures invariant “Structural Proxies” that provide a more efficient and physically grounded resolution of localization determinants, particularly in evolutionarily distant proteomes where primary sequence identity is low.

While these pLMs achieve impressive accuracy on benchmark datasets, they suffer from three critical limitations: (1) they function as “computational black boxes” [15], hindering mechanistic interpretation essential for biological hypothesis generation; (2) they may rely on evolutionary correlations learned from training data rather than universal biophysical principles, making their predictions difficult to interpret mechanistically; and (3) they fail to explicitly model the sequence-structure relationship central to Anfinsen’s dogma, where primary sequence determines three-dimensional structure [16]. Although graph representation learning offers a promising way to model protein structures [17], it is still constrained by the availability of high-quality 3D data.

Recent work has increasingly focused on incorporating biochemical knowledge into graph-based representations for protein-related tasks. One representative method is PSICHIC [18], which predicts binding affinities by constructing a bipartite interaction graph from protein sequences and ligand SMILES. PSICHIC is based on modeling intermolecular recognition and leverages predicted protein contact maps together with covalent ligand bonds to build a relational scaffold for simulating binding dynamics using a Graph Neural Network (GNN). In contrast, BioGraphX focuses on intramolecular structure and is driven by structural proxies. BioGraphX constructs a homogeneous constraint graph directly from the protein sequence. This construction relies on explicit biophysical rules, such as hydrophobic packing and salt bridges. Importantly, it does not depend on learned structural priors or external ligand information. While PSICHIC focuses on inter-molecular interactions, BioGraphX emphasizes the structural constraints encoded within a protein sequence that determine its cellular localization.

This work addresses three specific questions: (1) Can biophysically-grounded graph encodings bridge the sequence-structure gap without requiring 3D structures? (2) Do such encodings generalize better than pLMs to evolutionarily diverse proteins? (3) Can explicit biophysical rules enable interpretable predictions that reveal mechanistic insights? We introduce BioGraphX, a framework that constructs multi-scale interaction graphs directly from sequences using biochemical constraint rules as illustrated in Figure 1. In contrast to LocPro [13] that implement biochemical features in a post-hoc manner, training embeddings first and then appending features, the proposed framework implements integration natively at the encoding stage.

**Figure 1:**
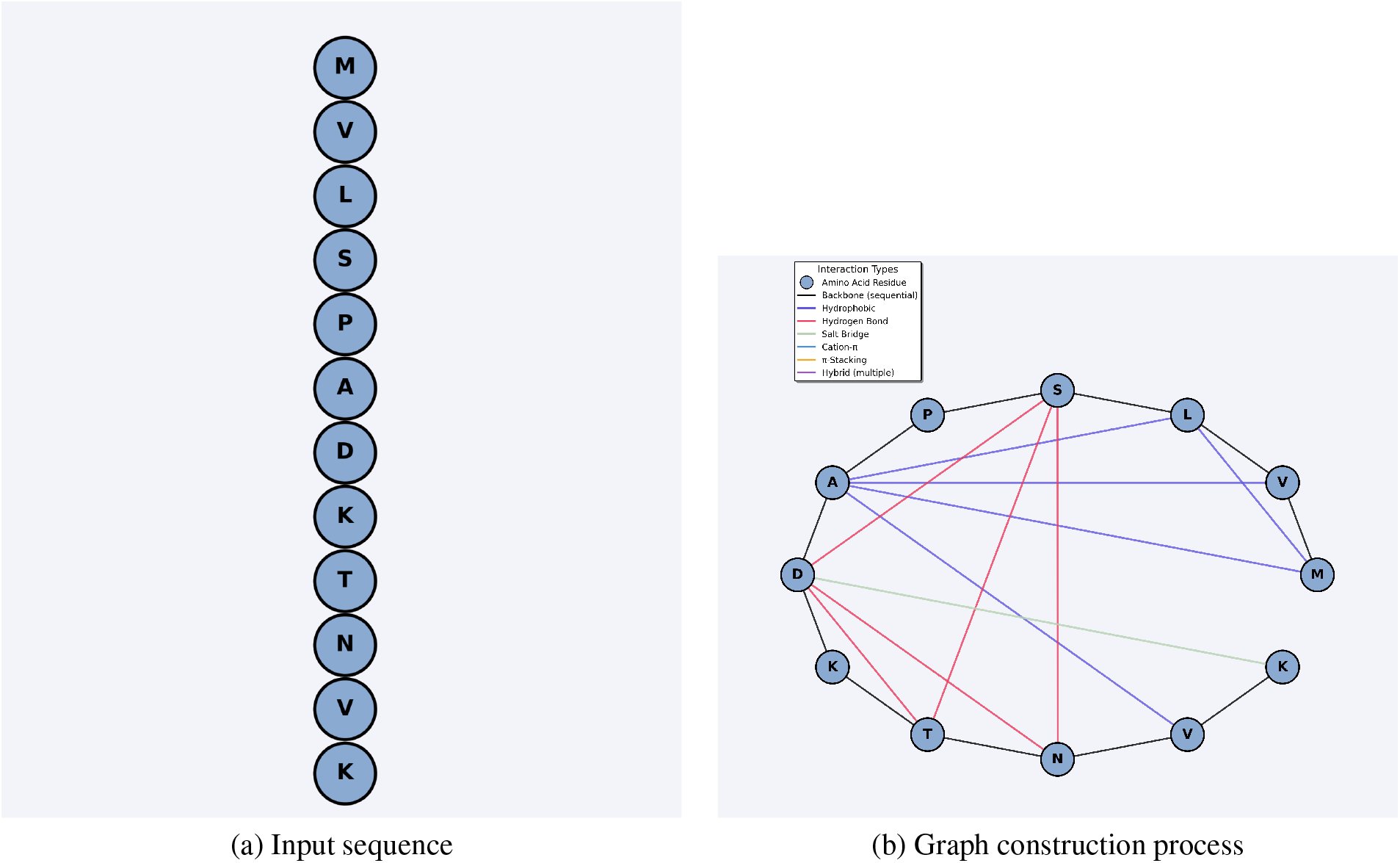
Detailed visualization of BioGraphX framework. (a) Input protein sequences for encoding. (b) Construction of protein residue interaction graphs with edges representing biochemical relationships.

BioGraphX-Net combines these encodings with ESM-2 embeddings through an interpretable gating mechanism that quantifies reliance on evolutionary vs. biophysical signals, enabling transparent decision-making. Unlike approaches that require end-to-end fine-tuning of the full language model (billions of parameters), our framework keeps the ESM-2 backbone frozen and trains only 13.46M specialized parametersa reduction of two orders of magnitude in trainable parameters compared to full fine-tuning, while maintaining competitive accuracy. This represents a knowledge-driven approach to model efficiency for proteome-scale prediction. Our contributions include: (1) the BioGraphX encoding algorithm; (2) the hybrid BioGraphX-Net architecture; (3) comprehensive benchmarking with superior generalization; and (4) explainable analysis revealing biophysical predictors of localization.

With its reusable, explainable, and structure-aware representation, BioGraphX takes an important step toward modeling the relationship between sequence, structure, and function.

## Methods

### Datasets and Preprocessing

This study relies on data from DeepLoc 2.0 [2] benchmark datasets. To ensure data integrity, a series of filtering steps was applied. The primary inclusion criterion required sequences to contain only the 20 standard amino acids. Accordingly, any entries with unknown or non-standard residues, such as ‘X’, were excluded from the analysis. Following this filtering process, 28,251 of the original 28,303 sequences in the DeepLoc 2.0 dataset met the inclusion criteria.

Evaluation followed the official DeepLoc 2.0 protocol using 5-fold cross-validation with a strict constraint of ≤30% sequence identity between training and test sets, predicting ten eukaryotic subcellular compartments (Cytoplasm, Nucleus, Extracellular, Cell Membrane, Mitochondrion, Plastid, Endoplasmic Reticulum, Lysosome/Vacuole, Golgi Apparatus and Peroxisome). This homology-reduced partitioning ensures that performance differences reflect model capability rather than sequence memorization.

To independently verify the structural proxy hypothesis, we employed the Escherichia coli eSol solubility benchmark according to the procedure outlined in the description of fGNNSol[19]. The database consists of experimentally determined solubility measurements of the proteome of Escherichia coli, whereby solubility is defined as the percentage of the supernatant fraction to the whole fraction obtained using PURE physicochemical experiments. The dataset was homology-reduced to retain only sequence pairs with global identity *<* 30% and E-value ≤ 1 × 10^−6^, yielding 2,679 protein sequences. We adopted the same partition as fGNNSol: 2,019 sequences for training, 268 for validation, and 392 for testing.

The choice of the above measure of solubility is because protein solubility is highly dependent on its three-dimensional structure, which encompasses the hydrophobic character of its surface, charge distribution, stability of folding, and aggregation properties, thus providing a good basis for assessing if constraints from sequence information are enough for obtaining structural meaning through graph modeling. Importantly, BioGraphX features were used without any modification from the localization task, and no ESM embeddings or deep learning components were employed in this validation, isolating the contribution of the graph physicochemical encoding alone.

### Overview of BioGraphX-Net Architecture

The BioGraphX-Net architecture is composed of three integrated stages. The first section, BioGraphX encoding, involves the construction of biochemically-calibrated constraint graphs from protein sequences to facilitate the extraction of multi-scale topological, knowledge-guided, and physicochemical features. Subsequently, ESM-2 sequence embedding is utilized to capture high-dimensional evolutionary context through attention-pooled residue representations. In the final stage, an interpretable gated fusion mechanism dynamically integrates physics-based and evolutionary signals via learned gating weights prior to multi-class classification. A schematic overview of this pipeline is presented in Figure 2.

**Figure 2:**
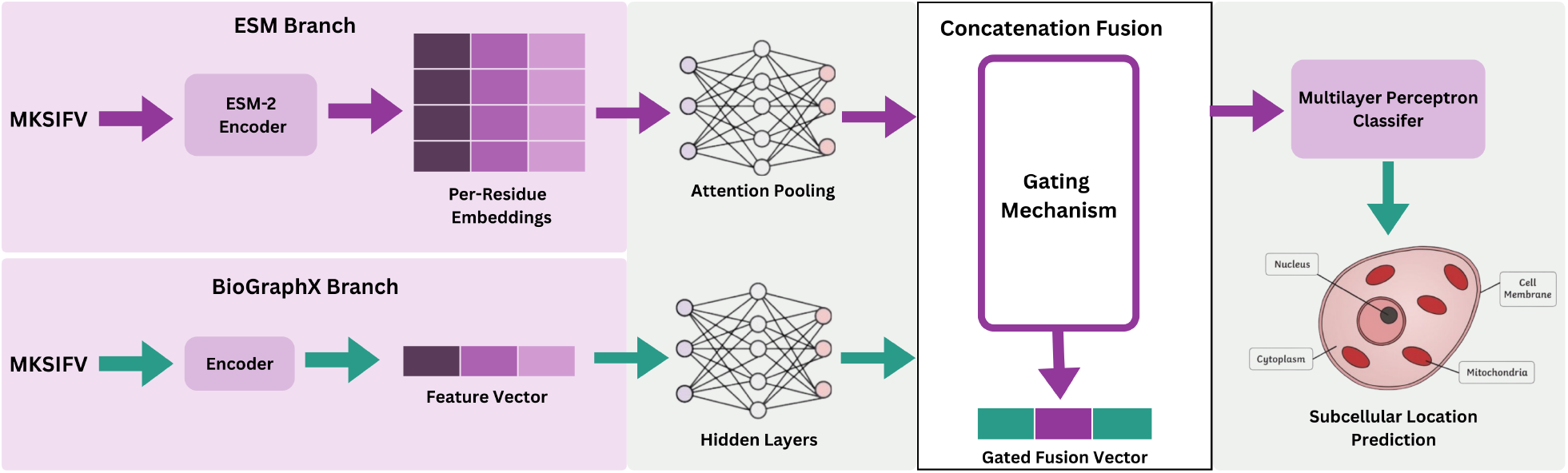
Schematic representation of the BioGraphX-Net computational pipeline. The diagram shows the integration of ESM sequence embeddings and BioGraphX physics-based features through a central gating mechanism, producing the combined Gated Fusion Vector for downstream classification.

### Sequence Embedding and Feature Extraction

#### ESM-2 Embeddings with Attention Pooling

The present study utilizes the ESM-2 model (esm2_t36_3B_UR50D) [10] to extract residue-level embeddings, providing a dense numerical representation for each sequence. To aggregate these variable-length embeddings into a fixed-length protein representation, an attention pooling mechanism was implemented [20]. This technique was employed to selectively weight the importance of specific residues, thereby ensuring that the most relevant biological signals for localization are captured. The raw ESM embeddings *E*_*seq*_ ∈ ℝ^*L×*2560^ are transformed into a weighted sequence representation *E*_*vec*_ according to the following operations:

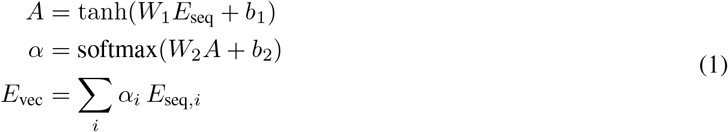

In this model, *A* stands for the attention transformation, while *α* refers to the masked attention weights that is assigned to each residue. After this aggregation step, the weighted representation, *E*_*vec*_, is passed through a bottleneck transformation. This step reduces its dimensionality and makes it easier to combine with other features. Specifically, the reduction to 1024 dimensions is performed by a fully connected layer, which is paired with Batch Normalisation (BN) and a ReLU (Rectified Linear Unit) activation function:

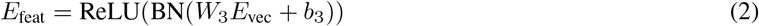

Here, *E*_vec_ is the original high-dimensional embedding produced by the ESM model. *W*_3_ and *b*_3_ are the learnable weight matrix and bias for the bottleneck layer. After this transformation, the resulting vector, *E*_feat_, represents the compressed semantic feature representation.

The purpose of this bottleneck reduction is to capture the most important features from the high-dimensional ESM embedding. This helps the model avoid overfitting to noise, while keeping the latent space compatible for later integration with biophysical features.

### BioGraphX Encoding Framework

To support proteome-scale analyses, the encoding pipeline provides parallel batch processing using Joblib, which significantly improves computational efficiency.

#### Foundational Biophysical Principles

The BioGraphX encoding framework converts protein sequences into multi-scale interaction graphs by applying biochemically calibrated constraint rules. While pLMs learn statistical patterns from large datasets, BioGraphX uses explicit physical laws supported by experimental data, resulting in deterministic and structure-informed features.

The framework relies on two main biophysical principles. The first, known as Anfinsen’s Principle, holds that a protein’s tertiary structure is determined by its amino acid sequence through encoded structural constraints [16]. BioGraphX implements these constraints explicitly via interaction rules between compatible residues, based on the understanding that specific residue pairs preferentially interact to stabilize fold topology. Secondly, within folded protein domains, residues that are proximal in the primary sequence are typically also proximal in three-dimensional space as a result of chain connectivity. This general consequence of polymer physics enables the construction of interaction graphs that serve as structural proxies without requiring explicit 3D coordinates. Maximum interaction distances were calibrated from the literature.

#### Graph Construction Algorithm

For a protein sequence *S* of length *L*, we construct an undirected weighted graph *G*(*V, E*) where vertices *V* = {*v*_1_, *v*_2_, …, *v*_*L*_} represent residues and edges *E* represent biochemical interactions. The complete algorithm is formalized in Algorithm 1.

##### Algorithm 1 BioGraphX Encoding Pipeline

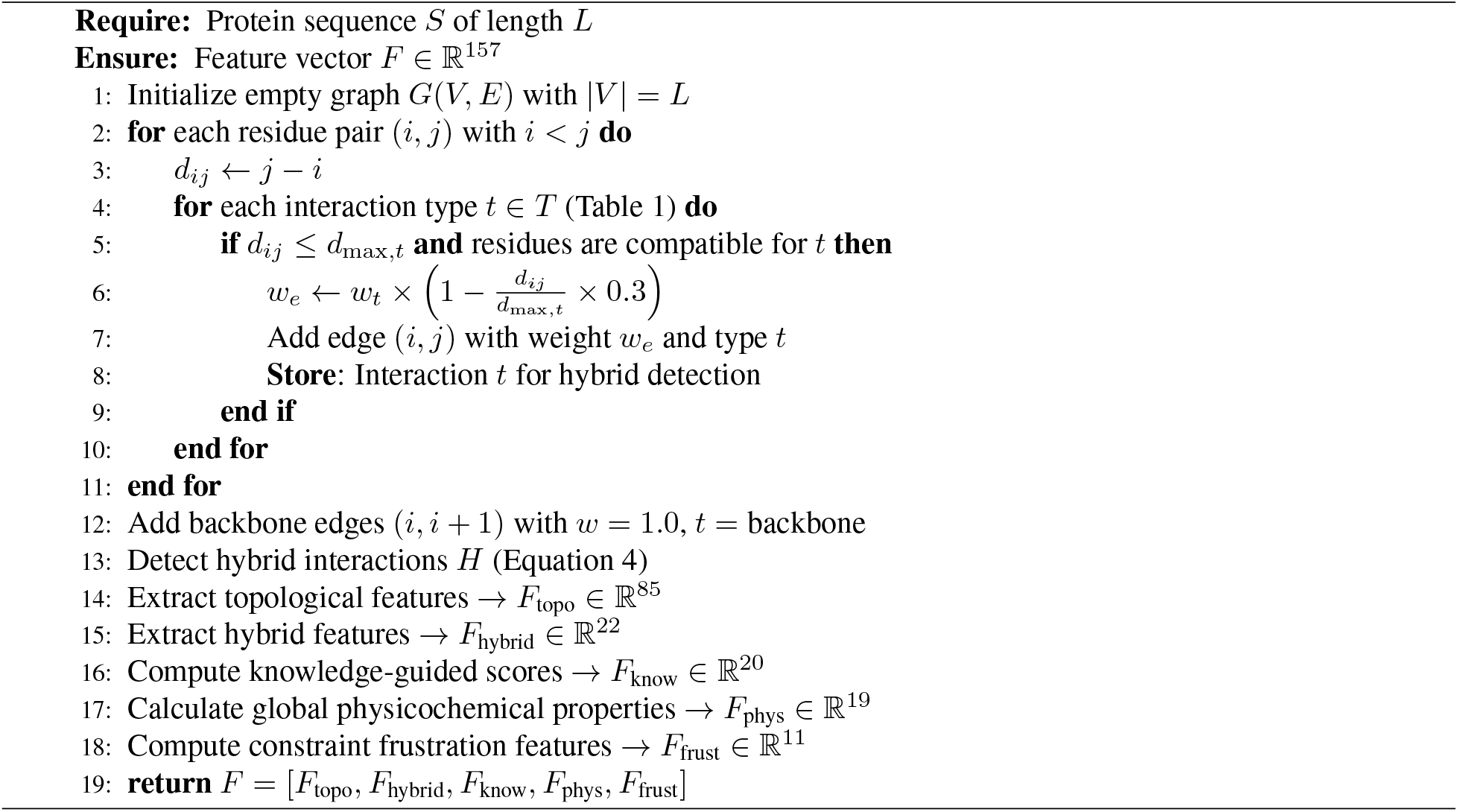

#### Biochemical Interaction Rules

The identification of the twelve interaction types and the calibration of their corresponding parameters were based on established biophysical literature and physical chemistry principles. It is important to clarify that all distance thresholds specified in this framework refer to linear sequence separation rather than three-dimensional spatial proximity. As Table 1 shows, these rules cover many types of forces, from the main stabilizing effects, like hydrophobic interactions [21, 28], to specific polar–aromatic motifs, including sulfur–*π* interactions [32]. The distance thresholds and the strength weights (*w*_*t*_) used by the graph enable the representation of the hierarchy of protein folding and stability using the protein sequence. This is achieved by using the distance thresholds to represent the maximum sequence distance over which these forces act to maintain structural integrity and the weights to represent the relative contribution of each type of interaction to biochemical stability. By using these deterministic rules, BioGraphX is able to generate a representation that is grounded in the principles of biophysics and acts as a robust proxy for 3D structural features.

**Table 1:** Biochemical Constraint Rules Used in BioGraphX.

| Constraint Type | Participating Residues | Strength | Max Dist | Key References |
| --- | --- | --- | --- | --- |
| Hydrophobic | A, V, L, I, M, F, W, Y, C | 1.0 | 20 | [21] |
| Hydrogen Bond | Donors: N, Q, S, T, Y, H, K, R, W<br>Acceptors: D, E, N, Q, S, T, Y | 0.8 | 12 | [22] |
| Salt Bridge | Positive: K, R, H<br>Negative: D, E | 0.7 | 35 | [23] |
| Disulfide Bond | C–C | 0.9 | 2000 | [24] |
| π-Interaction | Aromatic: F, Y, W | 0.6 | 15 | [25, 26] |
| Cation-π | Positive: K, R, H<br>Aromatic: F, Y, W | 0.65 | 10 | [27] |
| Van der Waals | A, V, L, I, M, F, W, Y, C, P | 0.4 | 6 | [28] |
| CH-π | Donors: A, V, L, I, M, C<br>Acceptors: F, Y, W | 0.3 | 8 | [29] |
| NH-π | Donors: N, Q, S, T, H, K, R, W<br>Acceptors: F, Y, W | 0.5 | 8 | [30] |
| Carbonyl–Carbonyl | D, E, N, Q, S, T, Y | 0.35 | 8 | [31] |
| Sulfur-π | Sulfur: M, C<br>Aromatic: F, Y, W | 0.45 | 8 | [32] |
| Backbone Interaction | Peptide backbone atoms | 1.0 | 1 | [33] |

In order to further enhance the representation, we have assigned a weight *w*_*t*_ ∈ [0, 1] to each type of interaction *t* and have proposed a distance decay function to account for the weakening of the influence of the interaction as the sequence distance increases. The weight *w*_*e*_ of a given edge *E* is given by:

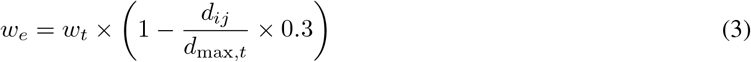

In this model, *d*_*ij*_ represents the sequence distance between two residues *i* and *j*, while *d*_*max,t*_ represents the maximum allowed separation for a particular type of interaction *t*. In addition, the constant factor of 0.3 is used to guarantee that a base level of interaction strength of approximately 70% remains even when the maximum threshold is reached. Results of ablation studies have shown that the choice of this factor has a significant impact: removing the decay reduced the accuracy by 1.44%, while increasing the factor to 0.5 leads to a 0.5% decrease in accuracy. This approach preserves a detectable signal for functional contacts despite increased linear separation, effectively balancing long-range constraint detection with noise reduction. This helps preserve functional contact signals across long sequence separations while mitigating noise. The weighting approach makes the graph sensitive to both the presence of a bond and how close the interacting residues are, providing a strong stand-in for structural features even without 3D coordinates.

#### Hybrid Interaction Detection

A key innovation of the framework is the detection of hybrid interactions, where two interaction types occur simultaneously between the same residue pair. These interactions serve as high-fidelity indicators of specialized structural motifs.

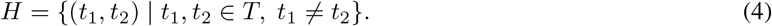

We define eight hybrid interaction types, assigning them increased contributions based on their biological significance, as summarized in Table 2.

**Table 2:** Hybrid Interaction Types.

| Hybrid Interaction | Biological Significance |
| --- | --- |
| SaltBridge-HBond | Common in enzyme active sites |
| Hydrophobic-Pi | Membrane protein anchoring |
| Cation-Pi-HBond | Nuclear localization signals |
| Hydrophobic-VDW | Hydrophobic core packing |
| Pi-Cation-HBond | DNA-binding interfaces |
| CH-Pi-Hydrophobic | Lipid bilayer interactions |
| Sulfur-Aromatic | Redox-active sites |
| Carbonyl-Charge | Electrostatic clusters |

**Table 3:** Feature Scales Used in BioGraphX.

| Feature Scale | # Features | Description | Key Metrics |
| --- | --- | --- | --- |
| Topological | 85 | Graph metrics quantifying network architecture | Node/edge counts, centrality, path metrics, modularity, communities |
| Hybrid | 22 | Co-occurrence patterns of interaction types | Salt-bridge / H-bond frequency, regional hybrid density |
| Targeting Signal Features | 20 | Features derived from known biological motifs | Regular expressions for NLS and targeting motifs |
| Physicochemical | 19 | Whole-protein physicochemical properties | Isoelectric point, GRAVY score, entropy |
| Constraint Frustration | 11 | Per-residue conflicting interaction energies | Signal–structure contrast, frustration localization, hotspot count |
| **Total** | **157** | **Multi-scale protein representation** | **Integrated sequence–structure features** |

**Note:** 81.5% of BioGraphX features (128/157) are graph-based and derived from biochemical interaction graphs rather than raw sequence patterns, providing structural proxies that complement evolutionary embeddings. **Detailed descriptions of all 157 features, including topological (Table S1), hybrid (Table S2), Targeting Signal Features (Table S3), physicochemical (Table S4), and frustration features (Table S5), are provided in the Supplementary Materials.**

#### Multi-Scale Feature Extraction

BioGraphX generates 157 features in five groups: topological, hybrid, knowledge-guided, global physicochemical, and constraint frustration. Topological features show network structure, hybrid features capture overlapping biochemical interactions, knowledge-guided scores use known targeting motifs, global physicochemical features describe general protein characteristics, and constraint frustration features quantify conflicting interaction energies. Hybrid interactions, in which two interaction types occur simultaneously between the same residue pair, are assigned weight multipliers according to their functional significance, as summarized in Table 2. This set of features relates to an approach based on the concept of frustration in protein folding [34], which focuses on the ways in which different energy types can interact in order to produce locally undesirable conformations. Frustration is calculated using the energy of interactions in the BioGraphX constraint graph (Algorithm 1). For each residue, we determine all edges associated with it and calculate the variance of their energies. If a residue undergoes the same type of interactions (hydrophobic interactions, for instance), then its frustration is minimal, whereas a residue undergoing two types of contradicting interactions (a salt bridge demanding a charge environment along with hydrophobic interactions demanding a non-polar environment) will show maximum frustration.

The complete set of extracted features, along with their definitions, is documented in Supplementary Material S1 **(see Table S1 for complete feature descriptions)**.

#### Sequence Adaptive Processing

Variable-length proteins were handled using a three-tier adaptive processing strategy. This method balances computational efficiency with preservation of biologically important information and is categorized by sequence length (*L*) as follows

- **Short sequences** (*L* ≤ 2, 000): These proteins were processed using their full sequences, as the computation required was considered negligible.
- **Medium sequences** (2, 000 *< L* ≤ 10, 000): The sequences were truncated smartly, keeping the first 40%, middle 20%, and last 40% of each sequence. The rationale for this specific split is that the N- and C-termini typically contain the most critical signalling and targeting motifs, respectively.
- **Long sequences** (*L >* 10, 000): A sliding window approach was employed. Feature aggregation for these sequences was carried out using an information content-weighted mechanism to prioritize high-entropy regions and ensure that functionally dense segments are represented.

The biochemical constraint graph encoding algorithm introduced in this study differs fundamentally from existing approaches in several key respects. Whereas traditional methods are generally limited to simple compositional features or require the acquisition of external structural data, contemporary approaches typically rely on “black-box” learned embeddings. BioGraphX distinguishes itself from these models by implementing explicit biochemical rules validated by experimental data.

### Hybrid Fusion Architecture

#### Dual-Branch Network Design

The hybrid architecture consists of two specialized input branches designed to process complementary protein representations. The first branch operates on ESM-2 embeddings, applying attention pooling followed by bottleneck compression to extract evolutionary sequence information.

The second branch processes BioGraphX features through an enhanced three-layer nonlinear transformation, substantially increasing representational capacity for modeling biophysical constraints. Formally, the BioGraphX transformation is defined as:

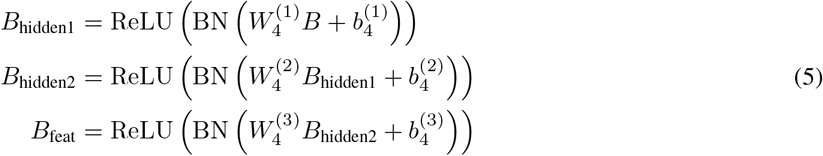

This expanded transformation projects the original 157-dimensional BioGraphX feature vector into a 1,024-dimensional latent space, matching the dimensionality of the ESM-2 embeddings. The successive nonlinear mappings enable the model to capture complex and higher-order relationships among biophysical constraints.

Finally, the outputs of the ESM and BioGraphX branches are fused using an adaptive gating mechanism (Figure 2), allowing the network to dynamically balance evolutionary and biophysical information during prediction.

#### Interpretable Gated Fusion Mechanism

The fusion stage employs an adaptive gating mechanism that dynamically balances the contributions of evolutionary embeddings and biophysical features on a per-protein basis. After obtaining the transformed feature representations *E*_feat_ ∈ ℝ^1024^ (Equation 2) and *B*_feat_ ∈ ℝ^1024^ (Equation 5), the two vectors are concatenated to form a joint representation:

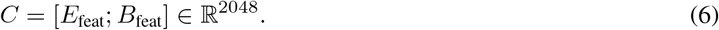

A two-layer gating controller then computes protein-specific weighting signals as follows:

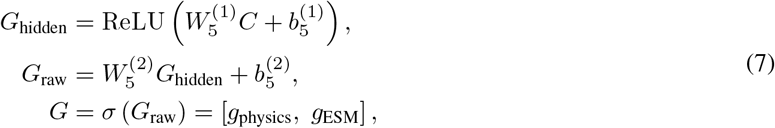

where *σ*(·) denotes the sigmoid activation function, producing normalized, per-protein gating coefficients. The gated feature vector is then computed via element-wise modulation of each feature block:

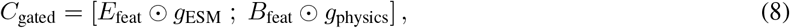

where ⊙ denotes element-wise multiplication, with each scalar gate value broadcast across the corresponding 1024-dimensional feature subspace. This formulation allows the model to independently regulate the influence of evolutionary and biophysical signals for each protein. The use of a two-layer gating controller provides greater expressive capacity than a single linear transformation, enabling the learning of complex and context-dependent fusion strategies.

#### Multi-layer Classifier

The last stage of classification consists of a multi-layer perceptron (MLP) which maps the gated features to the sub-cellular compartment labels. The architecture consists of a 1024-dimensional hidden layer with Batch Normalisation, followed by a 512-dimensional layer and an output layer representing the 10-class logits:

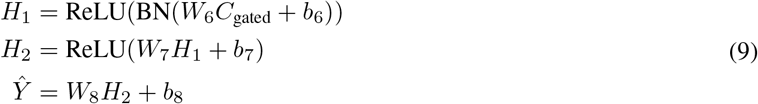

where *H*_1_ and *H*_2_ are hidden layer activations, and *Ŷ* is the final prediction. The resulting model contains approximately **13.46 million parameters**, achieving a balance between high representational capacity and computational efficiency.

### Training Protocol and Optimization

The classification thresholds were optimized by maximizing the Matthews Correlation Coefficient (MCC) [35] for each individual class. The optimization procedure employed a Focal Loss with parameters *α* = 0.25 and *γ* = 2.0 to address class imbalance and handle difficult examples. Optimization was carried out using the AdamW optimizer with a base learning rate of *η*_base_ = 1 ×10^−5^ and weight decay of 1 ×10^−2^, together with a cosine annealing learning rate scheduler. Differential learning rates were applied across model components: the physics branch used *η* = 3 ×10^−5^, the ESM branch used *η* = 1 × 10^−5^, the gating controller used *η* = 0.5 × 10^−5^, and the classifier used *η* = 1 × 10^−5^. During the early stages of training, a physics encouragement loss with coefficient *λ* = 0.1 was applied to promote balanced utilization of evolutionary and biophysical information.

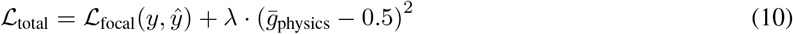

where ℒ_focal_ is the Focal Loss, 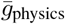 is the mean value of physics gate in a batch, and *λ* = 0.1 balances the usage of both information resources in early training stages. After epoch 10, the auxiliary term is removed (*λ* = 0).

Gradient clipping was applied in a component-specific manner, with a maximum norm of 5.0 for the physics branch and 1.0 for all remaining components.

The training protocol comprised the following procedures: (1) five-fold cross-validation was employed while maintaining ≤30% sequence identity between folds; (2) an early stopping strategy with a patience of 10 epochs was implemented; (3) a batch size of 64 was used; and (4) the maximum number of training epochs was set to 35. The final model was selected based on the highest observed validation performance.

### Explainability Framework

BioGraphX makes its predictions interpretable via four mechanisms that allow the researcher to analyze decisions at the level of each residue.

- **Gate Analysis:** Looking at the gating weights *G* makes it easy to see how much the various proteins use evolutionary versus biophysical features, by how much one is more prevalent.
- **SHAP Feature Importance:** Shap analysis [36] framework was applied to gain an insight on the contribution of biophysical features. SHAP values (*ϕ*_*ijk*_) for each protein *i*, feature *j*, and class *k* were calculated using a Kernel Explainer with stratified sampling in order to balance the representation from all subcellular compartments. Feature importance for each class was measured as average absolute SHAP value for all the proteins.

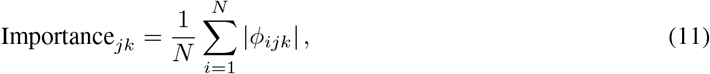

where *N* denotes the total number of proteins in the evaluation set. In order to isolate the contribution of biophysical features, while controlling for the information of evolutionary features, a special PhysicsWrapper was built into which ESM embeddings were fixed and BioGraphX features were allowed to vary. This design allows pinpointing localization choices made during development to very specific biophysical constraints, independent of evaluating evolutionary signal strength.
- **Knowledge-Guided Score:** The direct mapping of model properties to known targeting motifs, e.g. nuclear localization signals, mitochondrial targeting peptides, or signal peptides, allows hypothesis to be generated biologically about the mechanism of the predictions.
- **Hybrid Pattern Visualization:** Regions containing clusters of specific interaction patterns, such as SaltBridge –HBond motifs near active sites, are highlighted to show where key structural features play a crucial role.

### Evaluation Metrics

We assessed the model using standard multi-label metrics to measure how accurate and robust the framework is. The following parameters were used:

- **Accuracy:** This measures the percentage of cases where the predicted labels match the actual labels, providing a clear sense of the model’s accuracy.
- **Jaccard Index:**This compares the similarity of set of predictable and real labels, and assists in checking accuracy of secondary localizations.
- **Micro F1:** This measures the overall balance of precision and recall across all protein–label pairs.
- **Macro F1:** This measures the average per compartment so that even rare classes affect the total score fairly.
- **Matthews Correlation Coefficient (MCC):** This measures performance for each location so that class imbalance does not skew the results [35].

## Results

### Feature Group Ablation Analysis

To measure the relative contributions of graph-based features and sequence-based features to the localization task, an exhaustive ablation study was performed on the DeepLoc 2.0 dataset. Four scenarios were considered with the same XGBoost hyperparameters, as shown in Table 4.

**Table 4:** Ablation study using the XGBoost model.

| Config | Accuracy | Jaccard | Micro F1 | Macro F1 | MCC |
| --- | --- | --- | --- | --- | --- |
| Full BioGraphX | 0.3747 | 0.5264 | 0.6191 | 0.4746 | 0.5766 |
| Without Graph | 0.3332 | 0.4839 | 0.5814 | 0.4317 | 0.5362 |
| Graph Only | 0.3087 | 0.4458 | 0.5521 | 0.3560 | 0.5118 |
| Signal Only | 0.2955 | 0.4447 | 0.5413 | 0.3862 | 0.4908 |

By excluding the graph-based features but including only the sequence-based features, there was an observed decrease in Micro F1 of 0.038 (from 0.619 to 0.581), in the Jaccard index of 0.043 (from 0.526 to 0.484), and in the MCC of 0.040 (from 0.577 to 0.536). This reduction in each measure confirms the value that graph-based features provide for prediction performance.

We note that graph features alone (Micro F1: 0.552) underperform sequence-derived features alone (Micro F1: 0.581). This is due to complementarity, since each source contributes unique information. While graph topology describes interaction, packing, hydrophobicity, and community information, sequence-derived features describe evolutionary signals and physical-chemical properties of proteins. Nevertheless, the fusion of both types performs better than any single configuration (Micro F1 = 0.619), illustrating a central aspect of our design within BioGraphX.

The decrease in the Jaccard Index after the graph deletion (0.526 to 0.484) carries an interesting significance in the context of multilabel prediction, since the Jaccard Index measures the intersection between the predicted labels and true labels for proteins sharing localization sites. This suggests that graph-based features offer some special benefit for resolving cases where there are multiple simultaneous locations for a protein, with possible targeting constraints encoded by interaction topology.

### Benchmarking Against State-of-the-Art

BioGraphX-Net outperforms DeepLoc 2.0 and LocPro (2025) on key multi-label metrics (Table 5). It achieves a Micro-F1 of 0.78, higher than DeepLoc 2.0 (0.73) and LocPro (0.76). Higher Jaccard (0.72) and Macro-F1 (0.69) scores also show better handling of multi-label predictions.

**Table 5:** Performance Comparison Across Localization Prediction Methods.

| Metric | YLoc+ | DeepLoc 2.0 (ESM1b) | DeepLoc 2.0 (ProtT5) | LocPro | BioGraphX-Net (Ours) |
| --- | --- | --- | --- | --- | --- |
| **Prediction Type** | Multi | Multi | Multi | Multi | Multi |
| **Num. Labels** | 1.57 ± 0.02 | 1.27 ± 0.02 | 1.26 ± 0.02 | - | 1.52 ± 0.02 |
| **Accuracy** | 0.32 ± 0.02 | 0.53 ± 0.02 | 0.55 ± 0.02 | - | **0.55 ± 0.01** |
| **Jaccard** | 0.50 ± 0.01 | 0.68 ± 0.01 | 0.69 ± 0.01 | - | **0.72 ± 0.01** |
| **Micro F1** | 0.56 ± 0.01 | 0.72 ± 0.01 | 0.73 ± 0.01 | - | **0.78 ± 0.01** |
| **Macro F1** | 0.42 ± 0.01 | 0.64 ± 0.01 | 0.66 ± 0.01 | - | **0.69 ± 0.01** |
| **MCC per location (↑)** |  |  |  |  |  |
| Cytoplasm | 0.38 ± 0.02 | 0.61 ± 0.01 | 0.62 ± 0.01 | 0.612 ± 0.002 | **0.64 ± 0.01** |
| Nucleus | 0.42 ± 0.02 | 0.66 ± 0.02 | 0.69 ± 0.01 | **0.718 ± 0.002** | **0.71 ± 0.02** |
| Extracellular | 0.61 ± 0.05 | 0.85 ± 0.03 | 0.85 ± 0.04 | **0.892 ± 0.003** | 0.86 ± 0.03 |
| Cell membrane | 0.44 ± 0.02 | 0.64 ± 0.01 | 0.66 ± 0.01 | 0.662 ± 0.006 | **0.67 ± 0.01** |
| Mitochondrion | 0.47 ± 0.02 | 0.73 ± 0.03 | **0.76 ± 0.02** | 0.754 ± 0.003 | **0.76 ± 0.03** |
| Plastid | 0.72 ± 0.02 | 0.88 ± 0.01 | **0.90 ± 0.01** | 0.816 ± 0.003 | **0.90 ± 0.01** |
| Endoplasmic reticulum | 0.17 ± 0.04 | 0.52 ± 0.01 | 0.56 ± 0.03 | 0.533 ± 0.007 | **0.59 ± 0.02** |
| Lysosome / Vacuole | 0.07 ± 0.03 | 0.24 ± 0.03 | 0.28 ± 0.04 | **0.341 ± 0.027** | 0.33 ± 0.04 |
| Golgi apparatus | 0.11 ± 0.04 | 0.36 ± 0.06 | 0.34 ± 0.05 | 0.373 ± 0.003 | **0.43 ± 0.07** |
| Peroxisome | 0.05 ± 0.02 | 0.48 ± 0.05 | **0.56 ± 0.08** | 0.524 ± 0.017 | 0.54 ± 0.05 |

**Note:** Performance values are taken from the DeepLoc 2.0 publication [2], while results for LocPro are reported as presented in the LocPro study [13]. All models were evaluated on the DeepLoc 2.0 benchmark dataset under identical experimental conditions. Metrics marked with “-” for LocPro were not reported in the original LocPro study and therefore could not be included in the comparison. Bold values indicate the best performance.

**Table 6:**
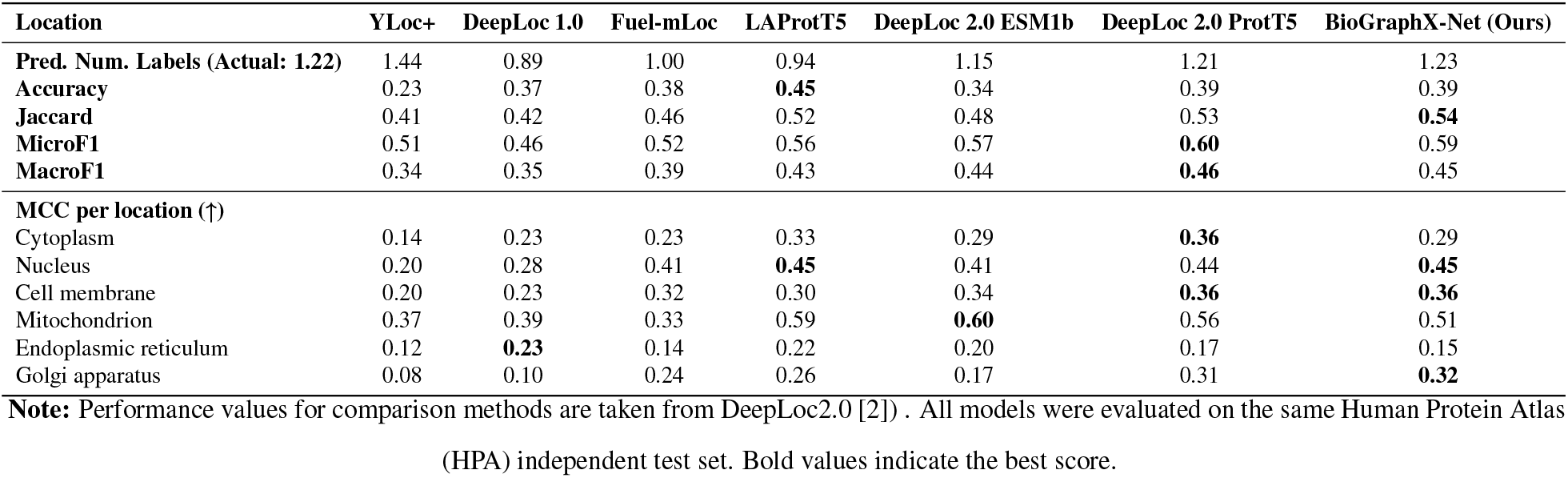
Performance comparison on the HPA independent test set.

While LocPro uses ESM-2 together with 1,484 sequence-based PROFEAT descriptors, BioGraphX-Net achieves similar overall performance and better results on hard organelles using just 157 graph features.

#### Organelle-Specific Performance and Sensitivity

A detailed per-location analysis of the Matthews Correlation Coefficient (MCC) highlights the distinct mechanistic advantages of structural graph encoding over sequence-only and expert-descriptor models. In particular, BioGraphX-Net shows superior resolution for organelles that are traditionally difficult to classify due to sparse data and dynamic interactions.

##### Challenging Compartments

For the Golgi apparatus, the framework achieved an MCC of 0.43, surpassing both DeepLoc 2.0 (0.34) and LocPro (0.373). Similarly, the model reached an MCC of 0.54 for the Peroxisome, which is higher than the 0.52 reported for LocPro and comparable to the 0.56 achieved by DeepLoc.

##### Targeting Signal Identification

In mitochondrial localization, BioGraphX-Net outperforms state-of-the-art methods, achieving an MCC of 0.76, surpassing the LocPro ESM2+PROFEAT hybrid (0.754) and matching DeepLoc 2.0 (0.76).

##### Standardized Benchmarks

Performance on the nucleus (0.71) and lysosome/vacuole (0.33) remains highly competitive on standardized DeepLoc benchmarks, outperforming DeepLoc 2.0 and matching LocPro.

Regarding multi-label prediction, BioGraphX-Net predicted an average of 1.52 labels per protein, exceeding the ground-truth average of 1.27 in the DeepLoc 2.0 dataset. While the higher Jaccard Index (0.72 vs. 0.69) and Micro F1 score (0.78 vs. 0.73) partly reflect improved recall, the elevated label count also indicates a tendency toward overprediction of secondary localizations compared to DeepLoc 2.0, which matches the ground-truth average of 1.27. This suggests that BioGraphX-Net’s higher sensitivity comes at the cost of reduced precision in multi-label assignment.

#### Generalization on Independent HPA Dataset

To evaluate generalization beyond the training distribution, BioGraphX-Net was benchmarked on an independent blind test set derived from the Human Protein Atlas (HPA; *N* = 1,717). This dataset provides a particularly rigorous testing environment because all included sequences share less than 30% similarity with the SwissProt training data. This is a critical challenge, as purely sequence-based models typically experience a sharp drop in performance under these conditions. Irrespective of this difficulty, BioGraphX-Net demonstrated a high level of robustness, as a Micro-F1 score of 0.59 was achieved with a conservative average of 1.23 predicted labels per protein.

We analyzed the gating mechanism and observed that the contribution pattern is protein-specific. BioGraphX features contributed 39.2%, while ESM embeddings contributed 50.5%. We further noted compartment-specific reliance: mitochondrial proteins depended approximately 40% on BioGraphX features, which is consistent with their physico-chemical properties. Conversely, Golgi apparatus proteins showed the lowest contribution from physics-based features and the strongest reliance on ESM embeddings, indicating a high degree of evolutionary conservation.

The adaptive nature of the gating ratios is further demonstrated by their distribution across proteins. We define balanced contribution as both gate values falling within 0.35–0.65 (neither branch contributing less than 35% or more than 65%), strong physics reliance as a physics gate exceeding 0.65 with ESM gate remaining below 0.65, and strong ESM reliance as an ESM gate exceeding 0.65 with physics gate remaining below 0.65. Under these definitions, 63.1% of proteins showed a balanced contribution between BioGraphX features and ESM embeddings, while 4.3% exhibited strong reliance on physics-based features and 32.6% exhibited strong reliance on ESM embeddings. The predominance of balanced contribution demonstrates that the gating mechanism genuinely integrates both information sources for the majority of predictions, rather than defaulting to one branch, confirming that evolutionary and biophysical signals provide complementary rather than redundant information at the protein level.

For example, nuclear proteins exhibited strong structural determinants, reflected by high physics contributions (up to 64.0%). In contrast, proteins with high ESM dependence (up to 79.8%) primarily relied on patterns of evolutionary conservation.

Notably, our model achieved superior performance on nucleus and Golgi apparatus with MCC of 0.45 and 0.32, respectively. The results suggest that BioGraphX-Net effectively captures physical principles that generalize across species.

Overall, these results suggest that explicit biophysical encoding can achieve performance comparable to that of significantly larger black box models.

#### Cross-Dataset Validation of the Structural Proxy Hypothesis

To validate that BioGraphX graph encodings capture genuine structural signal beyond the localization task, we evaluated the 157 physicochemical features, without any ESM embeddings and without the neural network architecture, on the *Escherichia coli* eSol protein solubility benchmark, a task where three-dimensional structure is a primary determinant of the target property. We followed the exact evaluation protocol of fGNNSol [19], a state-of-the-art solubility prediction method that employs AlphaFold3-derived 3D structural features including DSSP secondary structure assignments, dihedral and bond angle features, intranode atomic pair distances processed through radial basis functions, and edge distance and direction features, totaling approximately 640 structural dimensions derived from explicit 3D co-ordinates, alongside ESM-C language model embeddings (1,152 dimensions) and global physicochemical features (16 dimensions), all processed through a dual-stream Graph Neural Network architecture combining Graph Convolutional Networks and an improved Graph Attention Network. The total input dimensionality of fGNNSol is approximately 1,831 dimensions.

In contrast, BioGraphX features were extracted purely from protein sequences and evaluated using a simple XG-Boost regressor with no access to 3D coordinates, no protein language model, and no deep learning architecture. This represents an 11.6× reduction in input dimensionality and the complete elimination of structural and evolutionary information sources. Despite the considerable variation in terms of input data, BioGraphX was still a comparable tool to fGNNSol in the case of eSol, tested through five random seeds (refer to Table 7). In the binary solubility prediction with the threshold of 0.5, BioGraphX demonstrated a recall value of 0.7262 ± 0.0075, which is almost identical to the recall value of fGNNSol, 0.734 ± 0.020. Likewise, the area under the ROC curve (AUC) for BioGraphX was 0.843 and that of fGNNSol was 0.898. As seen from the regression measures, a certain degree of variance in terms of performance can be expected given the considerably less input data available to BioGraphX (*R*^2^ = 0.431 ± 0.004 vs. 0.578).Nevertheless, the highly competitive recall and AUC values demonstrate that BioGraphX is capable of identifying biologically meaningful boundary conditions for the protein solubility classification task while using approximately 11.6× fewer features and without relying on explicit 3D structural coordinates.

**Table 7:** Five-seed evaluation (XGBoost with BioGraphX features vs fGNNSol)

| Metric | Our Mean ± Std | fGNNSol Mean ± Std |
| --- | --- | --- |
| $R^2$ | $0.4308 \pm 0.0043$ | $0.578 \pm 0.012$ |
| Pearson | $0.6578 \pm 0.0029$ | $0.763 \pm 0.008$ |
| RMSE | $0.2363 \pm 0.0009$ | $0.207 \pm 0.003$ |
| ACC | $0.7566 \pm 0.0071$ | $0.785 \pm 0.017$ |
| Precision | $0.7119 \pm 0.0098$ | $0.812 \pm 0.024$ |
| Recall | $0.7262 \pm 0.0075$ | $0.734 \pm 0.020$ |
| F1 | $0.7189 \pm 0.0075$ | $0.771 \pm 0.018$ |
| AUC | $0.8428 \pm 0.0023$ | $0.898 \pm 0.008$ |
| MCC | $0.5045 \pm 0.0141$ | $0.589 \pm 0.034$ |

Feature importance analysis on this structure-dependent task from the best-performing XGBoost model (seed 2024, Recall = 0.7381) provided direct mechanistic validation of the structural proxy hypothesis. Triangle_Count, a purely topological graph metric, ranked as the single most important feature (importance: 0.0842). In BioGraphX, Triangle_Count counts three-residue cliques in which all pairs simultaneously satisfy biophysical interaction distance thresholds derived from established biochemical literature, hydrophobic contacts within 20 sequence positions, hydrogen bonds within 12, and van der Waals interactions within 6. This sequence-derived triangulation serves as a structural proxy for local residue packing density, capturing biochemically constrained interaction triplets that correlate with physical compactness, complementing the explicit 3D packing information that fGNNSol measures through AlphaFold3 atomic pair distances and contact-based graph construction. The emergence of Triangle_Count as the dominant predictor on a structure-dependent task, from a model with no access to 3D information, constitutes direct empirical evidence that BioGraphX graph topology functions as a structural proxy.

Hydrophobic cluster features ranked second and third (Mean_Hydrophobic_Cluster_Size, Max_Hydrophobic _Cluster). In BioGraphX, these metrics are computed by constructing a subgraph restricted to hydrophobic residues connected through the interaction graph and measuring connected component sizes, capturing the same hydrophobic arrangement information that fGNNSol derives from DSSP-identified *α*-helices and *β*-sheets exposing hydrophobic residues. Notably, five of the top ten features were graph-based, underscoring the importance of graph-derived structural proxies. The fGNNSol paper explicitly states that DSSP features reveal the structural basis of solubility by identifying arrangements where hydrophobic residue exposure increases aggregation risk, the identical biophysical signal captured by our sequence-derived hydrophobic cluster topology.

Charge and electrostatic features appeared at ranks 5, 6, 8, and 9 (Net_Charge, Isoelectric_Point, Charge_Density _Physics, Net_Charge_Density). fGNNSol’s interpretability analysis identified net charge per residue as the single most important among its 16 global physicochemical features. The independent convergence of both models on electrostatic features as primary physicochemical determinants, across two distinct tasks, two datasets, two model architectures, and two completely different input representations, provides strong cross-model validation that these features capture genuine biophysical signal rather than task-specific artifacts.

One of the key findings concerns the comparison of the two models’ feature importance rankings. Since the interpretability analysis conducted on *fGNNSol* was limited only to its 16 global physicochemical features, it did not allow measuring the significance of its structural dimensions based on AlphaFold3 due to the non-linearity involved in the multi-layer GNN aggregation process. This is why BioGraphX’s cross-task feature importance addresses this interpretability issue. The fact that the features related to graph topology (`Triangle_Count` ranked first) are followed by features corresponding to hydrophobic clusters (ranked 2–3) and preceded only by the `Net_Charge` feature indicates that the structural aspect related to the local packing density of residues (encoded via explicit atom positions in *fGNNSol*) plays a leading role in solubility. This positions BioGraphX as providing mechanistic interpretability that structure-based models cannot achieve through their own architectures.

This cross-task convergence, in both predictive performance on classification metrics and in the identity of the most important biophysical determinants across independent models, provides strong empirical evidence that BioGraphX physicochemical graph encodings function as effective structural proxies, capturing folding constraints and physico-chemical properties that generalize across distinct structure-dependent prediction tasks without requiring explicit 3D structural information.

### Explainability Analysis

#### Gating Mechanism: Evolutionary versus Biophysical Signal Weighting

Analysis of the gate distribution across 5,442 validation samples revealed that the BioGraphX-Net architecture employs a complex, adaptive gating mechanism that dynamically balances evolutionary and biophysical information. Overall, the model explains an average of 60.8% ± 9.9% of the variation through ESM embeddings and 39.2%± 9.5% through BioGraphX physics features, corresponding to a ratio of 1.55:1. Statistical testing confirms significant differential gating between the two information sources (*t* = 108.48, *p <* 0.0001). Notably, only a weak negative correlation between the two gates was observed (*r* = −0.084), indicating largely independent modulation rather than strict competition.

The architecture also shows specific use of biophysical attributes by organelles based on the complexity of biology. Organelles that have complex mechanisms of importing proteins and molecules, including the TOC/TIC complex in plastids, Pex5/Pex7 receptors in peroxisomes, and TOM/TIM complex in mitochondria, rely most heavily on physics-based attributes, owing to their requirements for proper target recognition through specific receptor complexes. In contrast, compartments where localization is primarily determined by well-conserved sequence motifs, such as the cell membrane (transmembrane helices), Golgi apparatus (short hydrophobic anchors), and lysosome/vacuole (dileucine and tyrosine-based sorting signals), demonstrated greater dependence on ESM embeddings.

These compartment-adaptive gating patterns suggest that BioGraphX-Net learns to modulate the degree of biophysical validation required for accurate localization, applying stronger physics-based reasoning where evolutionary similarity alone is insufficient and relying more heavily on sequence homology where it provides a reliable signal, as shown in Figure 3.

**Figure 3:**
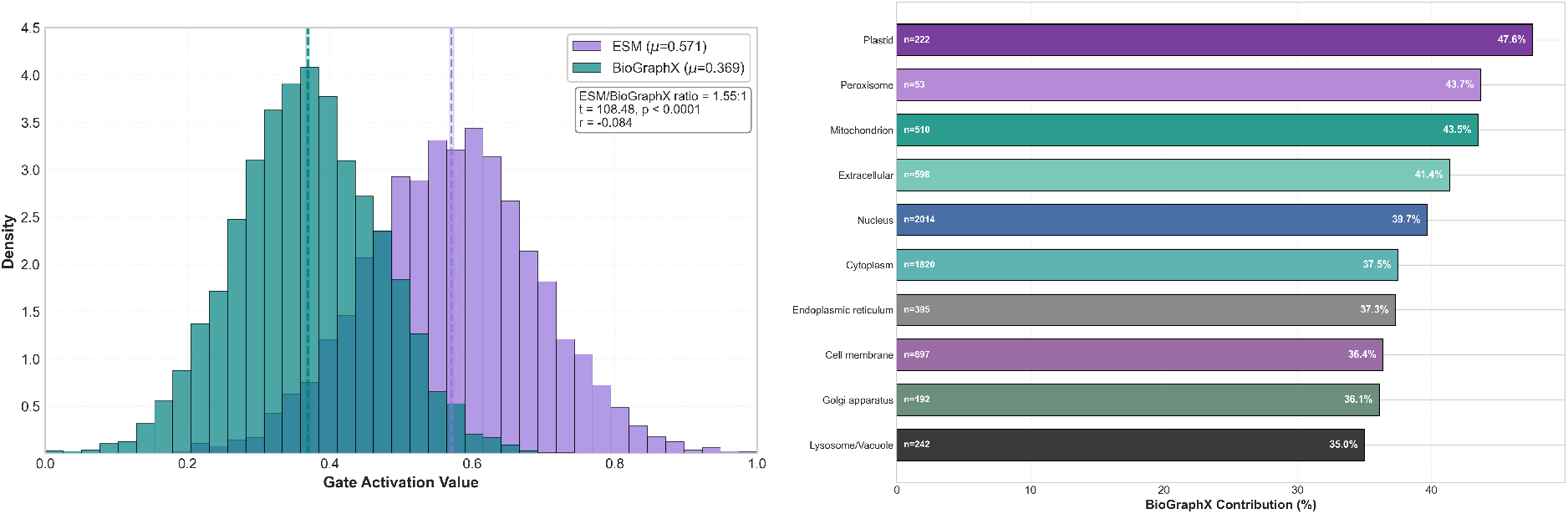
(A)Distribution of dynamic gating weights. The gating mechanism assigns consistently higher activation to sequence embeddings while maintaining a substantial contribution from BioGraphX features, demonstrating complementary feature fusion. (B)Quantitative measurement of reliance on BioGraphX features across organelles.

#### SHAP Analysis

SHAP values quantify model-internal feature importance, the contribution of each feature to the model’s output for a given prediction. They do not directly prove biological causation, nor do they establish that the model’s decision process mirrors biological mechanisms. All interpretations in this section describe computational patterns observed in feature importance analysis, contextualized with established cell biology literature.

To elucidate the biophysical rules underlying BioGraphX-Net predictions, we performed SHAP analysis on 200 stratified protein samples drawn from the validation partition. This approach quantifies the directional contribution of each of the 157 BioGraphX features across ten eukaryotic subcellular compartments, thereby revealing compartment-specific biophysical feature importance patterns. In addition to improving model interpretability, this analysis identifies which biochemical features the model prioritizes for each compartment, patterns that, when contextualized with established cell biology literature, suggest biophysical rules consistent with known localization mechanisms.The resulting ranking of physics-based feature utilization is illustrated in Figure 3B.

#### Directional Impact of Biophysical Features

The SHAP analysis showed a surprising pattern: the vast majority of features in the Signal are so-called repellers, not attractors (Figure 4). This finding suggests that the implementation of a biologically plausible exclusionary logic with localization being determined by what a protein is not rather than what it is, is brought about in BioGraphX-Net. The most important feature, Signal_Cell_membrane_Residue, had strong negative SHAP values of 7 of the compartments but positive contribution for cell membrane localization.

**Figure 4:**
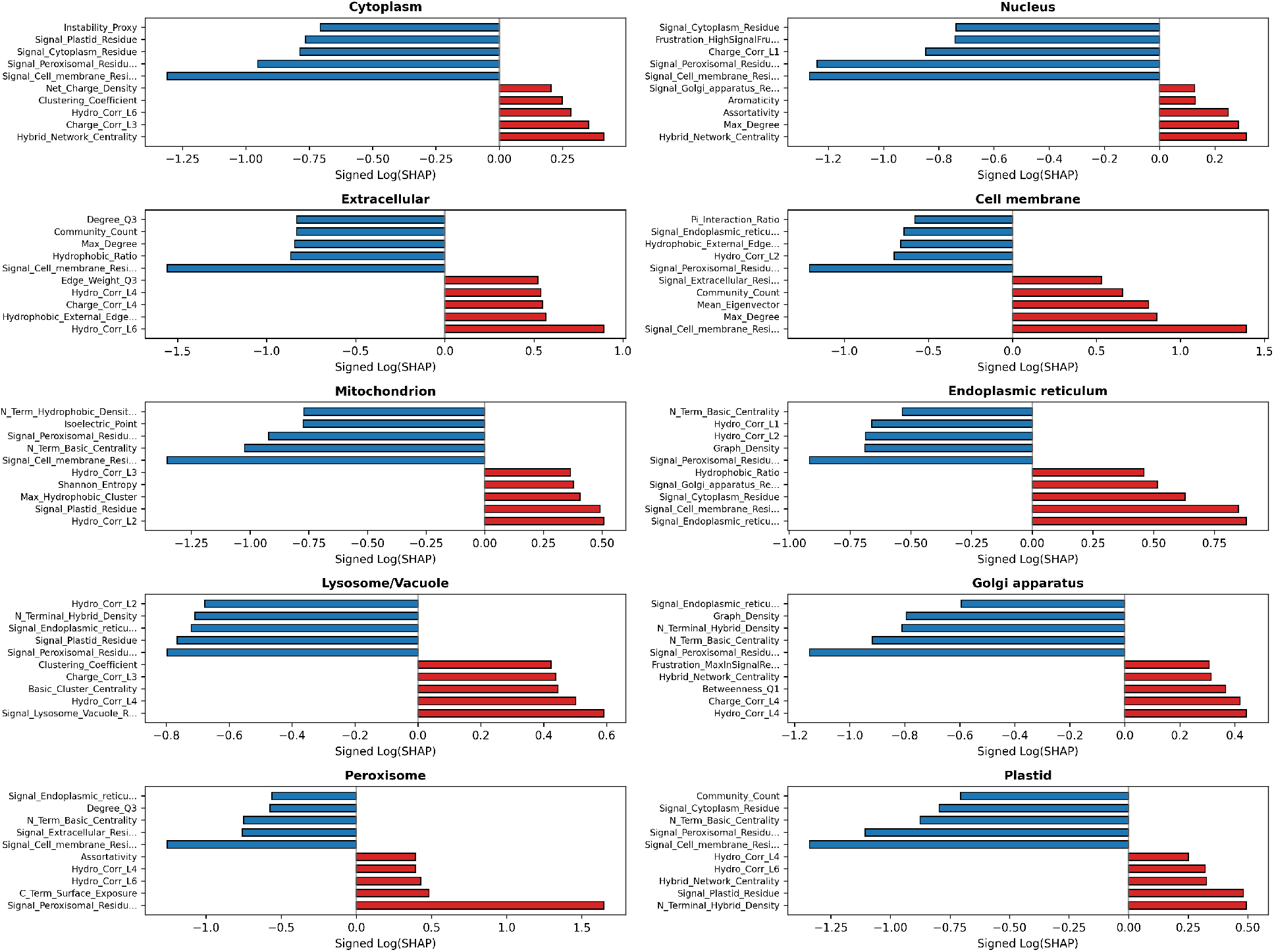
Top five organelle-specific features ranked by signed log-transformed feature importance values for improved visibility.

This is supported by the biology where membrane-bound proteins have unique physiochemical characteristics that inherently exclude them from soluble compartments. Likewise, Signal_Peroxisomal_Residue had high compartment specificity where the SHAP value was about 1.6 for Peroxisome, while it was negatively valued in all other compartments. This idea of exclusivity stems from the principle of biology where different organelles have unique signals through which only specific proteins can enter [37]. Nuclear localization uses nuclear pores [38], mitochondria use TIM/TOM complex [39], while peroxisomes use Pex5 receptor [40], making SHAPs detectable by BioGraphX Net.

#### Cross-Compartment Signal Affinities in Protein Targeting

SHAP analysis suggests counterintuitive cross-compartment signal affinities, providing insights into the functional architecture and evolutionary constraints of cellular trafficking systems. Convergent signal attractions are consistent with that organelles function as integrated hubs rather than isolated destinations.

In the following analysis, we operationally define several terms describing SHAP-derived relationships between compartments. Affinity refers to the SHAP value of a given Signal feature for a target compartment, where positive values indicate the feature’s contribution toward predicting that compartment. Attraction describes a positive SHAP value of one compartment’s Signal feature toward another compartment’s prediction (e.g., plastid Signal features showing positive SHAP for mitochondrion). Association refers to a co-occurrence pattern where Signal features from multiple source compartments show positive SHAP values for the same target compartment. Co-attraction describes reciprocal positive SHAP relationships between two compartments’ Signal features. Self-repulsion refers to negative SHAP values of a compartment’s own Signal feature for its own prediction, indicating that the feature acts as an exclusionary filter rather than a positive predictor for that compartment.

Notably, the Golgi, cytoplasm, and membrane signals show a strong association with the endoplasmic reticulum (ER), reflecting its central role in the biosynthesis of secretory and membrane proteins [41]. This pattern highlights the functional integration of the ER: all membrane proteins are synthesized in the ER [42], all secreted proteins are processed there [43], and both types of proteins subsequently transit to the Golgi apparatus [44, 45]. The ER and Golgi thus form a continuous secretory network, which may explain the co-attraction of their Signal features, as shown in Figure 4.

Similarly, the plastid Signal’s attraction to mitochondria is consistent with their shared endosymbiotic ancestry, capturing conserved targeting features in transit peptides that distinguish these organelles from non-endosymbiotic compartments [46, 47]. The cytoplasmic Signal shows self-repulsion, which may reflect cytoplasmic localization being determined more by the absence of competing signals than by the presence of specific motifs, an exclusion-based targeting logic.

The Golgi Signal’s attraction to nucleus predictions likely reflects shared intrinsically disordered regions, as both Golgi-resident glycosyltransferases and nuclear transcription factors contain polar/charged-enriched disordered tails with similar amino acid compositions [48, 49]. Frustration features may contribute the structural validation needed to differentiate true organelle-specific signals from compositional similarities.

Overall, these cross-compartment affinities suggest that BioGraphX-Net captures hierarchical targeting constraints and functional relationships within the cellular transport network, rather than relying on simple one-to-one sequence-to-compartment mappings. The model’s SHAP values suggest it treats organelles as integrated entities, emphasizing the endoplasmic reticulum (ER) as a key convergence point, the functional continuity between compartments of the secretory pathway, and the common targeting logic of organelles that have related functions.The model’s behavior suggests a complex representation of cellular organization that transcends simple cytoplasmic compartmentalization.

#### Frustration Features Resolve Targeting Ambiguities

The proposed features based on frustration also present compartment-specific properties (see Supplementary Table S5 for complete descriptions of frustration features). These features may also reflect the intricate import mechanisms of proteins. In nuclear localization, Frustration_HighSignalFrustration was identified as the fourth most important feature. In the endoplasmic reticulum, Frustration_CTerminal_Mean was also identified as a notable feature. In both cases, the negative SHAP values indicate that higher frustration decreases the predicted probability. This suggests that highly frustrated regions may create difficulties in the localization decisions of these compartments, as can be seen in Figure 4.

This also relates to the idea of frustration in protein folding, which states that interactions between residues can create conflicts in the energy of the protein sequence [50]. In the context of cellular compartments, these conflicts can relate to unclear signals that require additional regulatory mechanisms. For example, in nuclear localization, proteins with high N-terminal frustration can use chaperone-assisted transport or modification after translation to efficiently import [51].

#### Hydrophobicity Periodicity Encodes Compartment Identity

The SHAP analysis revealed that hydrophobic autocorrelation at certain sequence lags is a feature that is dependent on protein compartments. For instance, Hydro Corr L6, which is used to identify hydrophobic repeating patterns of 6 residues, was found to be the strongest positive predictor of extracellular proteins. The pattern of 3.6 residues is often seen as a helix [52]. In contrast, Hydro_Corr_L2 consistently contributed negatively to predictions for membrane, endoplasmic reticulum, and lysosomal compartments, indicating that short-range hydrophobic repetition is poorly tolerated in these targeting contexts.

#### Graph Topology Features Distinguish Structural Contexts

The relationship of Graph_Density with Golgi or endoplasmic reticulum predictions suggests that structural interaction patterns may influence localization to these compartments. Some enzymes, such as glycosyltransferases, that are located in the Golgi have one membrane anchor and a catalytic domain that is located in the lumen [49]. The number of strong interaction networks that enzymes such as this can have is restricted, which is why it is not good for graph density to be high for enzymes that are located in these compartments. The SHAP patterns suggest the model captures structural biases encoded in protein sequences. Also, larger hydrophobic patterns (lags of 4-6) could be associated with stable helical loops that facilitate proteins moving out of the cell and staying outside of it [53]. Shorter periodicities, on the other hand, may correspond to *β*-strand–like or structurally irregular regions, which appear more compatible with membrane-associated environments [54].

#### Charge Distribution Patterns as Targeting Determinants

N-terminal charge features exhibited pronounced compartment-specific effects. Notably, N_Term_Basic_Centrality shows strong negative SHAP values for mitochondrial, Golgi, and plastid predictions. This may seem strange at first because, of course, mitochondrial localization signals are well known to be amphipathic alpha helices with high basic residue content [55]. However, this may be explained when one recalls that this is a graph-based algorithm, and it is not just the basic residue content that is important, but the centrality of those basic residues to the entire network. In mitochondrial presequences, basic residues may be distributed along the periphery of the network rather than occupying highly central positions, which is consistent with the model down-weighting this feature despite overall charge enrichment.

Charge periodicity features provided further insight into localization-specific coding. Charge_Corr_L4 contributed positively to extracellular and Golgi predictions, suggesting that regularly spaced charged residues may support interactions with components of the secretory pathway or enhance solubility during transit through these compartments [56].

#### Compartment-Specific SHAP Patterns

The inclusion of all the SHAP value patterns leads to the identification of characteristic organelle-specific feature importance distributions that are organized according to polarity. While signal-based features contribute negatively to SHAP values in the vast majority of compartments, representing how the model learns to differentiate between compartments, organelle-specific sets of graph topology, hydrophobicity periodicity, charge distribution, and frustration patterns contribute positively in their corresponding compartments. Such patterns serve as descriptive representations of organelle-specific feature importance distributions based on SHAP values.

#### Evolutionary Insights from SHAP Patterns

Several SHAP patterns also point to underlying evolutionary signals. The strong negative contribution of Signal_Plastid _Residue to cytoplasmic predictions likely reflects the evolutionary separation between plastid-targeting signals, which originate from cyanobacterial endosymbionts, and the default localization landscape of the eukaryotic cytoplasm [46]. In a similar vein, the shared negative effect of N_Term_Basic_Centrality on both mitochondrial and plastid predictions suggests common constraints faced by import systems that evolved from endosymbiotic ancestors.

#### Context-Dependent Switch Features Enable Precise Discrimination

SHAP analysis revealed that BioGraphX-Net’s feature importance patterns are consistent with context-dependent ‘switch’ features that may allow the model to distinguish between evolutionarily convergent targeting signals. Rather than acting as simple attractors, these features show bidirectional SHAP patterns: positive contributions toward compatible compartments and negative contributions toward incompatible destinations, as shown in Figure 5.

**Figure 5:**
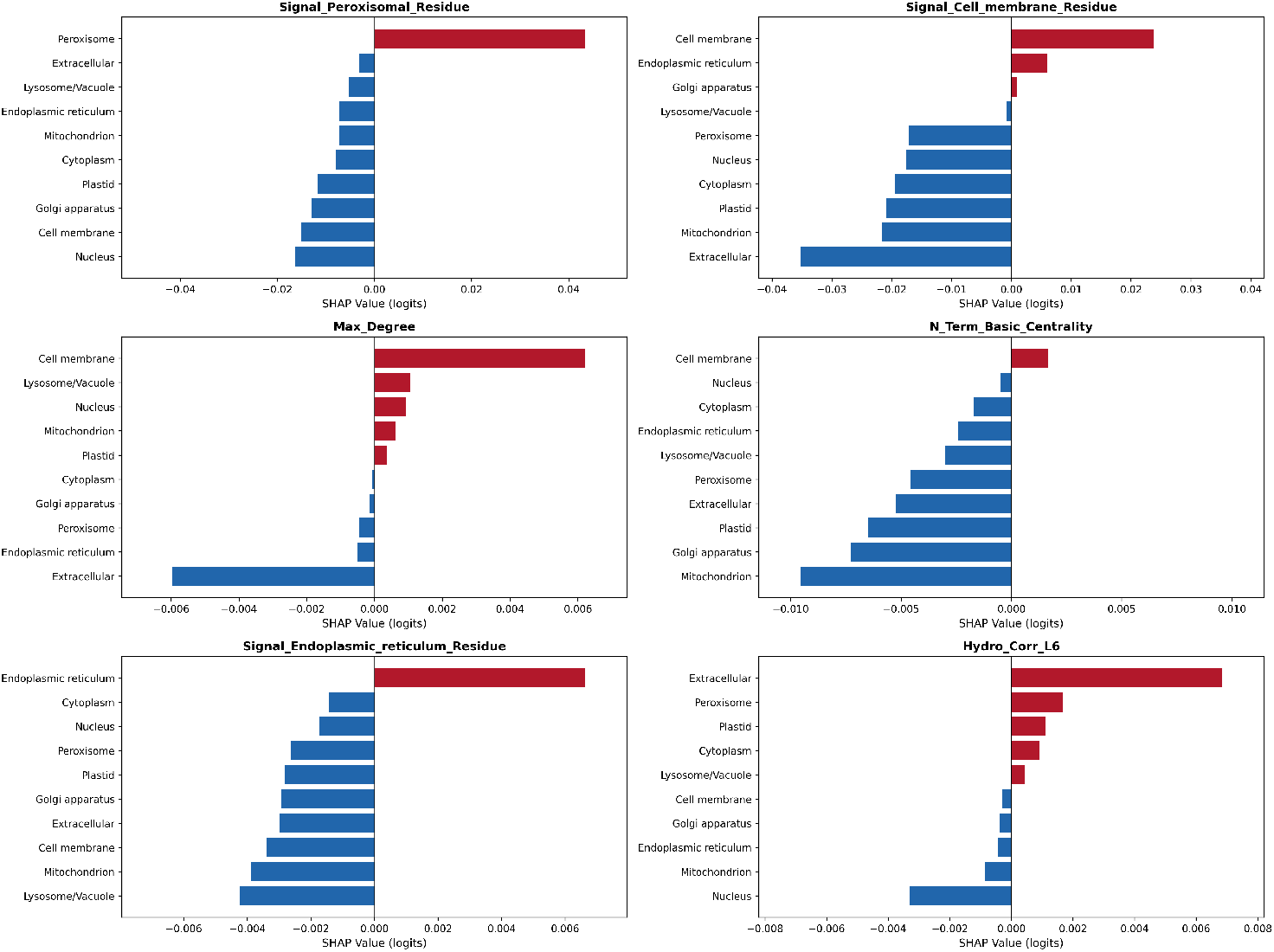
Bar chart highlighting six switch features that exert opposite SHAP impacts in different organelles.

This phenomenon can be observed through Hydro_Corr_L6, which shows a strong positive SHAP value for Extracellular proteins, consistent with the requirement for helical hydrophobic periodicity in secreted proteins.

By incorporating these conditional interactions, BioGraphX-Net’s SHAP patterns are consistent with addressing the phenomenon of evolutionary mimicry. Proteins destined for mitochondria and plastids retain targeting signals inherited from their respective endosymbiotic ancestors. Due to this shared evolutionary origin, these signals, such as N-terminal amphipathic helices with basic residues, can exhibit structural and physicochemical similarities that cause sequence-based models to confuse mitochondrial and plastid targeting. The model’s incorporation of biophysical features may help disambiguate these evolutionarily related signals. This pattern is analogous to cellular quality control mechanisms, though we note this analogy is descriptive rather than mechanistically proven. The positive correlation between Hydro_Corr_L6 and extracellular localization may also suggest optimization of these features during evolution. Proteins destined for the extracellular compartment require very specific hydrophobic motifs in order to traverse the secretory pathway efficiently without aggregating. The model’s SHAP values may reflect these requirements through the detection of long-range hydrophobic periodicity, which is consistent with what is currently known in the field of secretory biology[57].

#### Organelle-Specific Biophysical “Fingerprints”

In the radar plot of feature category contributions, each compartment shows a distinct pattern of biophysical feature importance (Figure 6). However, the patterns are not constant; they vary depending on the compartment. In most of the compartments, knowledge Signal features are dominant. This is a logical expectation, as motif recognition based on a sequence has to be the primary localization cue. In many cases, the first question the cell asks is: does this sequence contain the right localization signal? However, things get a bit more complicated when we take into consideration the sophisticated localization machineries of some other organelles. Frustration plays a unique role for the endoplasmic reticulum and the Golgi bodies. These localization machineries are part of the secretory pathway. And these localization signals are sometimes ambivalent and overlap.In these cases, the frustration feature may serve as a disambiguating signal that helps the model differentiate among these very similar signals and avoid misclassifications. But perhaps the most intriguing pattern of contributions is the one we find in the nucleus. In the nucleus, we find a nice balance of contributions by the knowledge signals and contributions by the topological and frustration features. This is consistent with the biological requirements of the nuclear localization machinery. In the nucleus, localization signals are not sufficient; structural compatibility to pass through the nuclear pores is a requirement as well.

**Figure 6:**
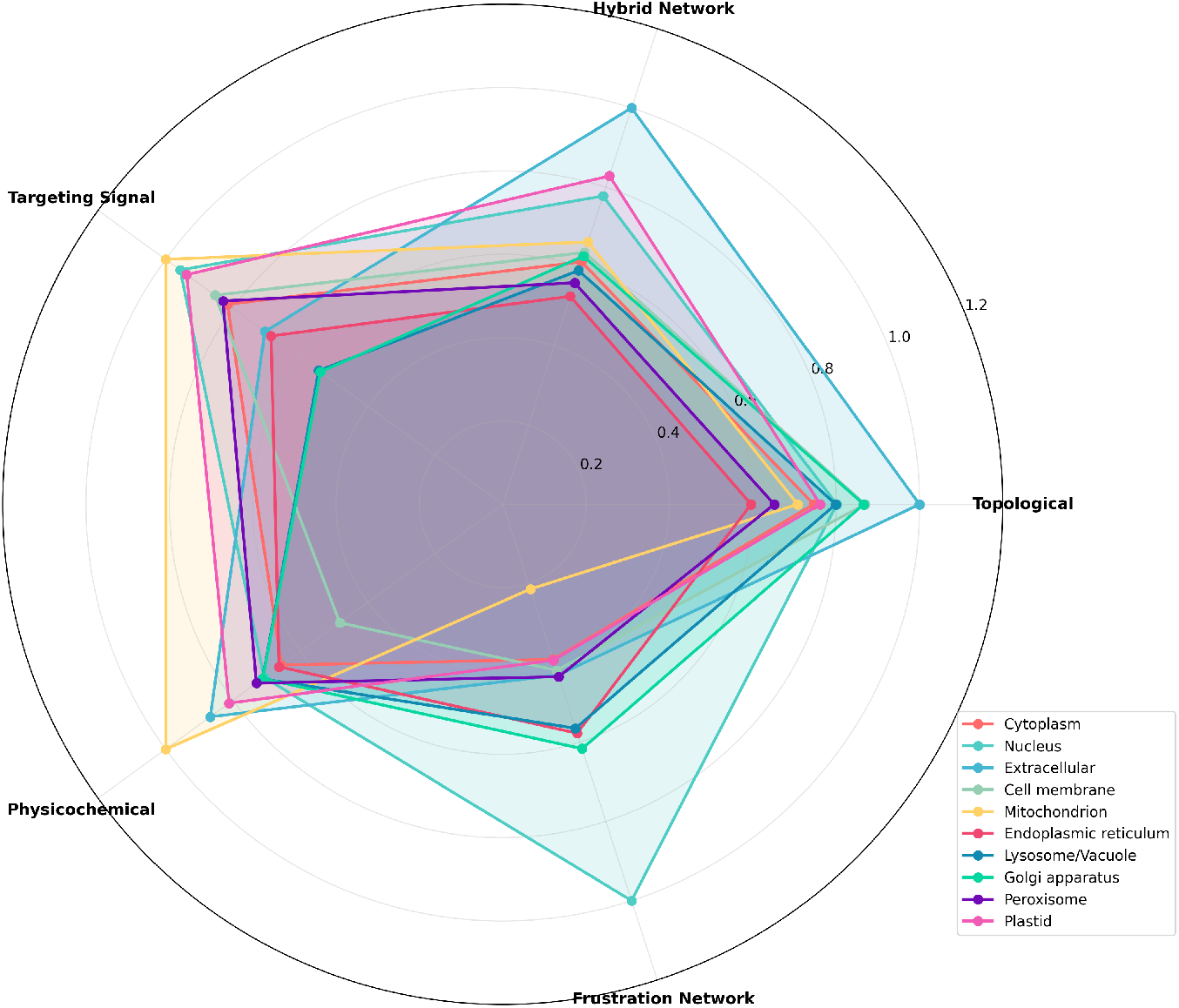
Radar chart summarizing the contribution of five feature categories.

In the above discussion, we have seen how the model’s SHAP patterns vary across organelles, and how these contribution patterns are consistent with the known biological localization machinery of the different organelles. In the plasma membrane and the cytoplasm, the knowledge Signal features are dominant, while in the nucleus, ER, and the Golgi apparatus,the topological and frustration features appear to complement the primary knowledge-guided signals. This suggests that the model’s feature importance patterns vary across compartments in ways consistent with the biological complexity of their localization machinery. Physicochemical features provide moderate, consistent contributions across compartments, acting as universal biophysical validators that complement the primary knowledge-based signals. Notably, the elevated frustration contributions in secretory compartments are consistent with cellular quality control mechanisms, where misfolded or mis-targeted proteins are actively rejected, while the balanced feature utilization in peroxisomes reflects their dual recognition system combining PTS signal detection with receptor-mediated structural compatibility.

## Discussion

### Toward Green AI in Proteomics

Beyond predictive accuracy, BioGraphX-Net demonstrates that explicit biochemical knowledge can achieve competitive performance without requiring large-scale computational resources. Like DeepLoc 2.0 and LocPro, our framework uses frozen ESM-2 embeddings, avoiding the computational cost of fine-tuning billion-parameter models. However, BioGraphX offers additional efficiency advantages: it captures structural proxies directly from sequence without requiring expensive 3D structure determination, and uses only 157 interpretable biophysical features compared to 1,484 PROFEAT descriptors in LocPro. The bottleneck compression from 2,560-dimensional ESM embeddings to a 1,024-dimensional latent space, combined with this compact feature set, results in 13.46M trainable parameters. Practically, this makes high-resolution localization predictions accessible on standard computational hardware, ESM inference is performed only once per sequence, and subsequent predictions require only the lightweight BioGraphX-Net architecture without GPU clusters. BioGraphX therefore moves from scale-focused AI to knowledge-driven AI, showing how encoding domain knowledge into compact, interpretable features can match the performance of more resource-intensive approaches in computational biology.

### Biological Implications: Capturing Complexity

What makes BioGraphX-Net stand out is its capability to grasp and visualize the real complexity of protein localization. The combination of evolutionarily relevant information and the biophysical laws of interaction makes it possible to predict the patterns of feature importance that reflect the complexity of residue-domain relationships and can easily be overlooked. Thus, BioGraphX-Net understands protein localization due to the intricacies of residue-domain relationships. SHAP patterns produced by the model indicate its ability to pick up both motifs and structural indicators. BioGraphX-Net is thus an embodiment of a crucial element of biological reality. Protein localization is not defined by sequence motifs but rather depends on their combination with the spatial organization and biophysics of interaction. All the results obtained show the significance of taking into account topological properties and biophysical interactions in computational modeling and the potential of bioinformatically generated hypotheses. Furthermore, they demonstrate the applicability of the proposed approach in case study analysis of proteins whose localization cannot be predicted based on the analysis of sequence motifs. One more insight provided by these studies concerns the mechanism of protein localization, namely that it depends not only on motifs, but also on their spatial organization.

### Cross-Task Validation Confirms Structural Proxy Hypothesis

The transfer of BioGraphX features to protein solubility prediction, a task where three-dimensional structure is the primary determinant of the target property, provides independent evidence that our constraint graph captures genuine structural constraints rather than merely statistical patterns in localization data. Using only 157 sequence-derived features and a simple XGBoost regressor, BioGraphX achieved recall comparable to fGNNSol (0.726 vs. 0.734), a state-of-the-art method that relies on explicit AlphaFold3 coordinates and a dual-stream graph neural network. The convergence of feature importance rankings, with *Triangle_Count*, hydrophobic cluster metrics, and net charge dominating both models, further demonstrates that BioGraphX encodes the same biophysical determinants that structure-based methods access through 3D data. This cross-task validation, combined with the strong performance on the independent HPA blind test, suggests that explicit biochemical rules can provide a generalizable structural proxy that complements evolutionary embeddings across diverse prediction tasks.

### Resolving the Eukaryotic Secretory Pathway

SHAP analysis produced an interesting result: features of the sequence Signal do not serve as simple attractors. Instead, the SHAP patterns suggest they may function as universal negative selection filters, a behavior that, to our knowledge, has not been previously characterized in an interpretable localization predictor

A good example of this is the localization of proteins to the Golgi apparatus. Initially, the sequence Signal of the Golgi apparatus was found to act as a negative predictor of localization to the Golgi apparatus itself, which produced an apparent contradiction or “self-repulsion. However, upon inclusion of the frustration features, the contradiction was resolved, suggesting a subtle two-tier validation mechanism. Thus, the sequence Signal of proteins, particularly those with similarities arising from disordered regions of proteins found in the nucleus or cytoplasm, was not sufficient, on its own, to result in a positive prediction of localization to the Golgi apparatus. It was necessary to provide an additional level of validation through biophysical means, such as hydrophobic edge patterns or sulfur-aromatic interactions, to result in a prediction of localization to the Golgi apparatus. This suggests that the sequence Signal alone was insufficient for prediction, but upon inclusion of additional validation, localization was correctly assigned. Thus, this computational behavior may reduce false positives arising from sequence mimicry, allowing genuine Golgi apparatus proteins to pass through structural validation while filtering out misleading similarities.

Importantly, this negative selection logic is not restricted to the Golgi. Membrane signals are highly repulsive to cytoplasmic and nuclear predictions. Peroxisomal signals negatively impact Golgi predictions. Finally, plastid-based predictions negatively impact cytoplasmic defaults. Based on SHAP patterns, we can see that the model considers not only the presence but also the absence of something. The use of frustration characteristics makes BioGraphX-Net able to include context information in addition to the Signal. Predictions cannot simply correspond to a motif but need to be backed up by biophysical evidence. This parallels quality control mechanisms within the cell, where targeting signals must be validated for structural compatibility before final decisions on localization are made.

The biological implications of this approach provide an explanation for how eukaryotic cells can accurately localize proteins while being composed of similar sequences, especially when comparisons are made to other compartments that are evolutionarily similar or structurally similars. Thus, BioGraphX-Net is more than just a pattern recognition algorithm. Its feature importance patterns mirror quality control logic analogous to cellular targeting validation mechanisms.

### Biological Significance of the Physics-Gated Fusion

The architecture of BioGraphX-Net is based on the observation that high-dimensional evolutionary embeddings and low-dimensional biophysical features are asymmetric. Placing these features side by side would unfairly favor evolutionary features because of their higher dimensionality. The higher dimensionality of the evolutionary features, together with their pretrained architecture, naturally gives them a dominant advantage in the learning process.

In order to achieve symmetry between the features, the network adds more representational power to the biophysical features through an intermediate hidden projection that expands the 157-dimensional BioGraphX vector to 1,024 dimensions, matching the ESM-2 embedding dimensionality. This enables the physics-based features to contribute more substantially alongside the evolutionary features during the gated fusion process. This architecture differs from simple post-hoc concatenation approaches, where biophysical features are appended to pre-computed embeddings without dedicated representational expansion, potentially limiting their influence on the final prediction.

Although the power of evolutionary features is substantial, they may sometimes be misleading. Sequence conservation may be retained for reasons apart from functional necessity, such as recent descent or neutrality, leading to false positives in which sequence identity is erroneously taken as evidence of similar localization..

By contrast, the biophysical features incorporated into BioGraphX allow for a direct consistency check. They encourage predictions that are consistent with physical and structural constraints, providing a complementary perspective to evolutionary signals. To reconcile these two forms of information, an adaptive gating mechanism is used, along with a physics-first training approach. This helps ensure that the biophysical information contributes meaningfully alongside evolutionary considerations. Instead of combining the features statically, the gating mechanism dynamically weighs the contribution of the evolutionary and biophysical signals for each protein. BioGraphX-Net thus produces a model that minimizes noise, avoids over-reliance on evolutionary considerations, and optimally balances the two forms of information. It does not privilege one view over the other but, rather, constantly balances between evolutionary memory and physical plausibility, just as the cell balances heritage and physical constraint.

### Comparison with Alternative Approaches

#### Graph Neural Networks on Protein Structures (GNN-3D)

The traditional Structure-Based Graph Neural Networks [17] require high quality three-dimensional protein structures, which can be obtained through AlphaFold2 [58] or through experiments using X-ray crystallography. These methods have been proven to perform well, especially for well-folded and structurally stable proteins. However, these methods are also accompanied by considerable limitations. First of all, the generation of structural predictions using AlphaFold2 for the entire proteome is computationally expensive. Second, predictions for intrinsically disordered regions are often low-confidence, which reduces reliability. Third, because these models depend on structure quality, errors in predicted coordinates can directly affect the learning results. BioGraphX bypasses these issues by learning approximate structures straight from amino acid sequences, so it doesn’t rely on full 3D coordinates.

Like PSICHIC [18], our approach favors knowledge-based graph encodings over black-box embeddings. But the two differ: PSICHIC uses chemistry-guided interactions to guide attention, while BioGraphX builds a physics-driven graph where the physicochemical rules define structure on their own. Graphs can either simulate processes like binding or directly represent structural properties like folding constraints. BioGraphX takes the second approach, using a sequence-based graph to capture structural determinants of localization, effectively applying Anfinsen’s postulate in a predictive and explainable way.

#### Contact Map–Based Methods

Contact prediction approaches such as PSICOV and MetaPSICOV [59] determine contact patterns between residues based on amino acid sequences alone, without resorting to structural data obtained experimentally, consistent with this basic principle behind BioGraphX, although applied for a different purpose downstream. While PSICOV and related contact predictors rely on coevolutionary statistics from MSAs [60], the BioGraphX approach relies on deterministic rules of biophysical interactions based on physical chemistry. The difference in these paradigms results in real-world differences in applicability, as MSA-based methods cannot be applied reliably to evolutionarily isolated proteins where MSAs cannot be built sufficiently deep, while the rule-based BioGraphX approach is independent of alignment depth.

#### Hybrid Feature Fusion Approaches

The most directly comparable hybrid localization method, LocPro combines protein language model (pLM) representations with external biological features through post-hoc feature concatenation. In such architectures, the learned pLM embeddings frequently dominate the decision-making process, resulting in feature asymmetry and limited interpretability of auxiliary signals. BioGraphX fundamentally departs from this paradigm by incorporating physics-based structural descriptors as first-class features within the model architecture. Through an interpretable gated fusion mechanism, balanced contributions from both evolutionary and biophysical sources are explicitly enforced. This architectural design accounts for the ability of BioGraphX to match the predictive performance of billion-parameter models while employing substantially smaller and computationally efficient networks.

### Universality of the BioGraphX Representation

Although the current implementation of BioGraphX is primarily concerned with the eukaryotic proteome, the underlying physicochemical graph encoding logic appears to be inherently domain-agnostic. The framework’s reliance on node-aligned topological descriptors suggests a natural extensibility to other biological macromolecules, such as RNA and DNA. By redefining node identities as nucleotides and edge constraints as base-pairing or stacking interactions, it is possible that BioGraphX could provide a unified ‘Structural Proxy’ for the entire Central Dogma. Specifically, the topological constants identified in this study, may represent universal physical requirements for the stability and localization of any linear biopolymer. Future research should be undertaken to explore the deployment of BioGraphX in predicting mRNA half-life and lncRNA subcellular trafficking, thereby positioning this framework as a universal sequence-to-physics encoder for the broader ‘omics’ landscape.

### Limitations and Future Work

BioGraphX-Net works well, but it has some limitations. Currently, it assumes that residues close in sequence are also close in 3D space. This usually works for folded domains, but it can miss long-range contacts. Future versions will use predicted contact maps from AlphaFold2 to better capture complex interactions.

Another limitation is that BioGraphX-Net is designed mainly for single-chain proteins. In complex systems, many proteins act as part of multimeric assemblies, and their subcellular localization is influenced by inter-chain contacts and overall quaternary geometry.

## Conclusion

In this study, we have introduced BioGraphX, a novel framework that bridges the sequence–structure gap by transforming primary protein sequences into multi-scale interaction graphs. By explicitly modeling biophysical constraints based on Anfinsen’s Dogma, the framework provides a deterministic constraint-based structural proxy that reduces reliance on high-quality 3D data. Our analysis results indicate that BioGraphX-Net outperforms DeepLoc 2.0 and LocPro on the DeepLoc 2.0 benchmark on challenging-to-predict organelles.

One of the main insights of our work is the standalone prediction ability of the biophysical representation. Using the DeepLoc 2.0 benchmark, we observe that a straightforward XGBoost model that uses just the 157 BioGraphX features, without any deep learning and ESM representations, obtains a Micro F1 score of 0.619 and MCC of 0.577 (see Table 4), which indicates that the biochemically-inspired rules can capture the localization patterns. Also, we validate the effectiveness of the same features by predicting the protein solubility task; we find that the recall rate reaches 0.726, very close to that of the current state-of-the-art approach fGNNSol (0.734) (see Table 7), which explicitly employs 3D structural information obtained from AlphaFold3 and deep graph neural networks.

SHAP analysis reveals feature importance patterns consistent with a universal negative selection strategy. Instead of acting as simple attractors, sequence signals appear to function as exclusionary filters that quickly rule out unlikely compartments, while organelle-specific combinations of graph topology, hydrophobicity periodicity, and frustration features guide precise localization. This two-tier system may help resolve evolutionary mimicry, such as the shared secretory signatures of the Golgi and ER, while preventing mislocalization through structural validation. This demonstrates that the framework can effectively combine evolutionary information with structural proxies in a transparent way.

BioGraphX pioneers a new paradigm of knowledge-integrated artificial intelligence for bioinformatics. By combining frozen language model embeddings with trainable biophysical graph features, we match state-of-the-art accuracy while training approximately two orders of magnitude fewer parameters compared to full fine-tuning of the ESM-2 model. This result demonstrates that encoding domain knowledge directly into model architecture is a powerful alternative to parameter scaling. The framework further provides mechanistic interpretation through built-in explainability tools and enables practical deployment, contributing to a more sustainable and interpretable future for machine learning in protein science.

## Acknowledgements

The author acknowledges the use of publicly accessible computational resources provided through the Kaggle platform for conducting the experiments in this study.

## Author contributions statement

A.S. conceived the study, developed the methodology, implemented the algorithms, conducted experiments, analyzed the data, and wrote the manuscript.

## Data Availability

The datasets analyzed in this study are publicly available. The DeepLoc benchmark dataset can be accessed from its original publication [2]. The complete BioGraphX prediction pipeline, including the encoding framework, ESM embedding generation, trained model weights, and the inference.py script for subcellular localization prediction, is available at: https://github.com/Abubakar-Saeed/BioGraphX.

## Supporting information

The following files are available. The following supplementary materials are available for this article:

- **Supplementary Material S1 (Table S1-S5)**: Complete BioGraphX Feature Descriptions - Contains comprehensive documentation of all 157 features across five categories:
  – Table S1: Topological Graph Features (85 features)
  – Table S2: Hybrid Interaction Features (22 features)
  – Table S3: Knowledge-Guided Signal Features (20 features)
  – Table S4: Global Physicochemical Features (19 features)
  – Table S5: Constraint Frustration Features (11 features)
- **Supplementary Material S2 (Table S6)**: Feature Summary and SHAP Framework - Overview of feature categories and explainability methods
- **Supplementary Material S3**: Computational Details - Time complexity and optimization strategies
- **Supplementary Material S4**: Parallel Implementation and Scalability - Details on parallel processing architecture

## Declaration of AI Use

During the preparation of this work, the author used Gemini in order to assist with manuscript formatting, language refinement, and organization of references. After using this tool/service, the author reviewed and edited the content as needed and take full responsibility for the content of the published article.

