## Supplementary Material for "BioGraphX: Bridging the Sequence-Structure Gap via Physicochemical Graph Encoding for Interpretable Subcellular Localization Prediction"

#### Supplementary Material S1: Complete BioGraphX Feature Descriptions (157 Features)

This supplementary document provides a comprehensive description of all 157 features extracted by the BioGraphX framework. The features are derived from a biochemically-calibrated constraint graph constructed directly from protein sequences. The framework's multi-scale approach captures protein topology, interaction networks, physicochemical properties, and constraint frustration to create a structural proxy without requiring 3D coordinates Gligorijević et al. (2021).

##### 0.1 1. Topological Graph Features (85 features)

*Description: Network architecture metrics derived from the biochemical interaction graph, applying concepts from graph theory to model protein residue relationships.*

Table S1: Topological Graph Features (85 total)

| Feature Name | Description | Biological Interpretation / Basis |
|---|---|---|
| Node_Count | Number of residues (vertices) in the graph. | Protein length. |
| Edge_Count | Number of biochemical interactions (edges) in the graph. | Overall connectivity. |
| Graph_Density | Ratio of actual edges to possible edges. | Compactness of interaction network. |
| Mean_Degree | Average number of interactions per residue. | Average residue connectivity. |
| Degree_Std | Standard deviation of degree distribution. | Heterogeneity in local connectivity. |
| Max_Degree | Maximum number of interactions for any residue. | Identification of highly connected hubs. |
| Degree_Q1 | First quartile of degree distribution. | Lower bound of connectivity. |
| Degree_Median | Median of degree distribution. | Central tendency of connectivity. |
| Degree_Q3 | Third quartile of degree distribution. | Upper bound of connectivity. |
| Mean_Weighted_Degree | Average edge-weighted degree (by strength). | Strength of interactions per residue. |
| Weighted_Degree_Std | Variability in weighted degree. | Diversity in interaction strengths. |
| Max_Weighted_Degree | Maximum weighted degree. | Residue with strongest total interactions. |
| Hydrophobic_Ratio | Proportion of hydrophobic interaction edges. | Dominance of hydrophobic packing. |

| Feature Name | Description | Biological Interpretation / Basis |
|---|---|---|
| HydrogenBond_Ratio | Proportion of hydrogen bond edges. | Importance of polar interactions. |
| SaltBridge_Ratio | Proportion of salt bridge edges. | Electrostatic stabilization strength. |
| Pi_Interaction_Ratio | Proportion of $\pi$-interaction edges. | Aromatic stacking interactions. |
| Disulfide_Ratio | Proportion of disulfide bond edges. | Extracellular/oxidized environment. |
| CationPi_Ratio | Proportion of cation-$\pi$interaction edges. | Charged-aromatic interactions. |
| VanDerWaals_Ratio | Proportion of van der Waals interaction edges. | Close-packing interactions. |
| CH_Pi_Ratio | Proportion of CH-$\pi$interaction edges. | Weak hydrophobic-aromatic interactions Nishio et al. (2014). |
| NH_Pi_Ratio | Proportion of NH-$\pi$interaction edges. | Polar-aromatic interactions. |
| CarbonylCarbonyl_Ratio | Proportion of carbonyl-carbonyl edges. | Backbone-backbone hydrogen bonding. |
| SulfurPi_Ratio | Proportion of sulfur-$\pi$edges. | Methionine/aromatic interactions Valley et al. (2012). |
| Backbone_Ratio | Proportion of backbone peptide bonds. | Chain connectivity and flexibility. |
| Mean_Betweenness | Average betweenness centrality. | Importance in information flow Brandes (2001). |
| Betweenness_Std | Standard deviation of betweenness. | Heterogeneity in pathway importance. |
| Max_Betweenness | Maximum betweenness centrality. | Critical bottleneck residues. |
| Min_Betweenness | Minimum betweenness centrality. | Peripheral residues. |
| Betweenness_Q1 | First quartile of betweenness. | Lower bound of centrality. |
| Betweenness_Q3 | Third quartile of betweenness. | Upper bound of centrality. |
| Mean_Closeness | Average closeness centrality. | Average proximity to all residues. |
| Closeness_Std | Standard deviation of closeness. | Heterogeneity in proximity. |
| Max_Closeness | Maximum closeness centrality. | Centrally located residues. |
| Min_Closeness | Minimum closeness centrality. | Remotely located residues. |
| Mean_Eigenvector | Average eigenvector centrality. | Influence in network neighborhoods. |
| Eigenvector_Std | Standard deviation of eigenvector. | Heterogeneity in neighborhood influence. |
| Community_Count | Number of detected communities. | Domain/module organization. |
| Modularity | Quality of community partition. | Strength of domain separation. |
| Mean_Community_Size | Average community size. | Typical domain size. |
| Community_Size_Std | Standard deviation of community sizes. | Heterogeneity in domain sizes. |
| Max_Community_Size | Largest community size. | Major structural domain. |
| Min_Community_Size | Smallest community size. | Minor structural element. |
| IntraCommunity_Edge_Ratio | Ratio of edges within communities. | Internal domain connectivity. |
| InterCommunity_Edge_Ratio | Ratio of edges between communities. | Inter-domain connectivity. |
| N_Term_Hydrophobic_Density | Hydrophobic density in N-terminal (1-30). | Signal peptide h-region indicator. |
| N_Term_Basic_Centrality | Centrality of basic residues in N-terminal. | Targeting signal importance. |
| N_Term_Clustering | Clustering coefficient in N-terminal. | Local compactness near start. |
| N_Term_Edge_Density | Edge density in N-terminal region. | Structural compactness near start. |
| C_Term_Surface_Exposure | Average degree in C-terminal (last 10). | Surface accessibility at terminus. |
| C_Term_Charged_Connectivity | Connectivity between charged in C-terminal. | Retention signal interactions. |
| C_Term_Terminal_Degree | Degree of the very last residue. | Terminal residue exposure. |
| Basic_Residue_Connectivity | Connectivity among basic (K,R,H) residues. | NLS cluster formation. |
| Basic_Cluster_Centrality | Centrality of basic residue clusters. | Importance of basic regions. |
| Basic_Cluster_Count | Number of connected basic clusters. | Multiple basic regions. |

| Feature Name | Description | Biological Interpretation / Basis |
|---|---|---|
| Max_Basic_Cluster_Size | Size of largest basic cluster. | Major basic region. |
| Hydrophobic_Cluster_ Connectivity | Connectivity among hydrophobic residues. | Hydrophobic core formation. |
| Hydrophobic_External_Edges | Edges from hydrophobic to non-hydrophobic. | Surface exposure of hydrophobic. |
| Hydrophobic_Cluster_Count | Number of hydrophobic clusters. | Multiple hydrophobic patches. |
| Max_Hydrophobic_Cluster | Size of largest hydrophobic cluster. | Major hydrophobic core. |
| Mean_Hydrophobic_Cluster_ Size | Average hydrophobic cluster size. | Typical hydrophobic patch size. |
| N_Term_Basic_Density | Basic residue density in N-terminal. | Mitochondrial/chloroplast targeting. |
| Basic_Gradient | Difference in basic density N- vs C-terminal. | Charge asymmetry. |
| Acidic_Gradient | Difference in acidic density N- vs C-terminal. | Acidic residue distribution. |
| Net_Charge_Density | Net charge per residue (KRH minus DE). | Overall electrostatic character. |
| Avg_Shortest_Path | Average shortest path length (LCC). | Global communication efficiency. |
| Graph_Diameter | Longest shortest path (LCC). | Maximum communication distance. |
| Global_Efficiency | Inverse of average shortest path length. | Network communication efficiency. |
| Local_Efficiency | Average efficiency of local neighborhoods. | Local communication robustness Brandes (2001). |
| Path_Length_Std | Standard deviation of sampled paths. | Heterogeneity in distances. |
| Max_Path_Length | Maximum sampled shortest path. | Extreme communication distance. |
| Clustering_Coefficient | Average local clustering coefficient. | Local cohesiveness. |
| Assortativity | Degree correlation coefficient. | Homophily in connectivity. |
| Graph_Radius | Minimum eccentricity. | Compactness measure. |
| Triangle_Count | Number of 3-node cliques. | Local motif density. |
| N_Region_Density | Edge density in first 30 residues. | N-terminal structural compactness. |
| M_Region_Density | Edge density in middle region. | Internal structural compactness. |
| C_Region_Density | Edge density in last 30 residues. | C-terminal structural compactness. |
| Mean_Edge_Weight | Average edge weight. | Average interaction strength. |
| Edge_Weight_Std | Standard deviation of edge weights. | Heterogeneity in strengths. |
| Max_Edge_Weight | Maximum edge weight. | Strongest individual interaction. |
| Min_Edge_Weight | Minimum edge weight. | Weakest individual interaction. |
| Edge_Weight_Q1 | First quartile of edge weights. | Lower bound of strengths. |
| Edge_Weight_Median | Median edge weight. | Typical interaction strength. |
| Edge_Weight_Q3 | Third quartile of edge weights. | Upper bound of strengths. |
| Edge_Weight_Range | Difference between max and min weights. | Span of interaction strengths. |

##### 0.2 2. Hybrid Interaction Features (22 features)

*Description: Co-occurrence patterns of biochemical interactions where two interaction types occur simultaneously between the same residue pair.*

Table S2: Hybrid Interaction Features (23 total)

| Feature Name | Description | Biological Interpretation |
|---|---|---|
| SaltBridge_HBond_Hybrid | Frequency of salt bridge + hydrogen bond. | Enzyme active sites, charged polar motifs. |
| Hydrophobic_Pi_Hybrid | Frequency of hydrophobic + $\pi$-interaction. | Membrane protein anchoring. |

| Feature Name | Description | Biological Interpretation |
|---|---|---|
| CationPi_HBond_Hybrid | Frequency of cation-π+ hydrogen bond. | Nuclear localization signals. |
| Hydrophobic_VDW_Hybrid | Frequency of hydrophobic + van der Waals. | Hydrophobic core packing. |
| Pi_Cation_HBond_Hybrid | Frequency of π-interaction + hydrogen bond. | DNA/RNA binding interfaces. |
| CH_Pi_Hydrophobic_Hybrid | Frequency of CH-π+ hydrophobic. | Lipid bilayer interactions. |
| Sulfur_Aromatic_Hybrid | Frequency of sulfur-π+ aromatic. | Redox-active sites. |
| Carbonyl_Charge_Hybrid | Frequency of carbonyl-carbonyl + salt bridge. | Electrostatic clusters, active sites. |
| N_Terminal_Hybrid_Density | Hybrid density in N-terminal region (1-30). | Complex targeting signals. |
| C_Terminal_Hybrid_Density | Hybrid density in C-terminal region (last 10). | Complex retention signals. |
| Hydrophobic_Region_Hybrid_Density | Hybrid density in hydrophobic clusters. | Specialized hydrophobic motifs. |
| Basic_Region_Hybrid_Density | Hybrid density in basic residue clusters. | Specialized nuclear targeting. |
| Hybrid_Edge_Ratio | Proportion of edges that are hybrid. | Overall interaction complexity. |
| Hybrid_Subgraph_Density | Density of subgraph of hybrid edges. | Compactness of hybrid clusters. |
| Hybrid_Network_Path_Length | Average path in hybrid subgraph. | Efficiency of hybrid network. |
| Hybrid_Cluster_Count | Connected components in hybrid subgraph. | Modularity of hybrid interactions. |
| Hybrid_Cooccurrence_1 | Highest co-occurrence score. | Most synergistic interaction pair. |
| Hybrid_Cooccurrence_2 | Second highest co-occurrence score. | Secondary synergistic pair. |
| Hybrid_Cooccurrence_3 | Third highest co-occurrence score. | Tertiary synergistic pair. |
| Hybrid_Diversity | Shannon entropy of hybrid types. | Diversity of hybrid interactions. |
| Hybrid_Network_Centrality | Average betweenness in hybrid subgraph. | Importance in hybrid network. |
| Hybrid_Clustering | Clustering coefficient of hybrid subgraph. | Local cohesiveness of hybrids. |

#### 0.3 3. Targeting Signal Features (20 features)

*Description: Sequence-based localization signals derived from known biological motifs and hybrid scores, informed by databases of experimentally validated targeting signals Negi et al. (2015); Nielsen (2017).*

Table S3: Knowledge-Guided Signal Features (20 total)

| Feature Name | Description | Biological Interpretation |
|---|---|---|
| Signal_Nucleus_Residue | Residue-based NLS motif score. | Presence of monopartite/bipartite NLS. |
| Signal_Nucleus_Hybrid | Hybrid-based nuclear compatibility. | Structural compatibility with nucleus. |
| Signal_Mitochondrion_Residue | Residue-based MTS score. | Amphipathic helix and charge patterns. |
| Signal_Mitochondrion_Hybrid | Hybrid-based mitochondrial compatibility. | Compatibility with mitochondrial import. |
| Signal_Extracellular_Residue | Signal peptide grammar score. | n-region, h-region, c-region patterns. |
| Signal_Extracellular_Hybrid | Hybrid-based extracellular compatibility. | Structural compatibility with secretion. |
| Signal_Cell_membrane_Residue | Transmembrane domain score. | Hydrophobic runs for membrane insertion. |
| Signal_Cell_membrane_Hybrid | Hybrid-based membrane compatibility. | Compatibility with membrane environment. |

| Feature Name | Description | Biological Interpretation |
|---|---|---|
| Signal_Endoplasmic_reticulum_Residue | ER retention signal score. | KDEL/HDEL/KKXX motifs. |
| Signal_Endoplasmic_reticulum_Hybrid | Hybrid-based ER compatibility. | Compatibility with ER lumen/membrane. |
| Signal_Golgi_apparatus_Residue | Golgi targeting score. | Tyrosine motifs (YXXΦ), TMD properties. |
| Signal_Golgi_apparatus_Hybrid | Hybrid-based Golgi compatibility. | Structural compatibility with Golgi. |
| Signal_Lysosome_Vacuole_Residue | Lysosomal targeting score. | Dileucine motifs, glycosylation signals. |
| Signal_Lysosome_Vacuole_Hybrid | Hybrid-based lysosomal compatibility. | Compatibility with lysosomal environment. |
| Signal_Peroxisomal_Residue | Peroxisomal targeting score. | PTS1 (SKL) and PTS2 motifs. |
| Signal_Peroxisomal_Hybrid | Hybrid-based peroxisomal compatibility. | Compatibility with peroxisomal import. |
| Signal_Plastid_Residue | Plastid transit peptide score. | Ser/Thr-rich, low acidic N-terminal. |
| Signal_Plastid_Hybrid | Hybrid-based plastid compatibility. | Compatibility with chloroplast import. |
| Signal_Cytoplasm_Residue | Cytoplasmic default score. | Absence of strong targeting signals. |
| Signal_Cytoplasm_Hybrid | Hybrid-based cytoplasmic compatibility. | Compatibility with cytoplasmic environment. |

#### 0.4 4. Global Physicochemical Features (19 features)

*Description: Whole-protein biophysical properties derived from amino acid composition and sequence Kyte and Doolittle (1982); Leuenberger et al. (2017), reflecting constraints important for protein evolution and stability Sikosek and Chan (2014); Rockah-Shmuel et al. (2015).*

Table S4: Global Physicochemical Features (19 total)

| Feature Name | Description | Biological Interpretation |
|---|---|---|
| Isoelectric_Point | Theoretical pI via bisection method. | Charge at physiological pH; solubility. |
| Net_Charge | Total charge at pH 7.0. | Overall electrostatic properties. |
| Charge_Density_Physics | Net charge per residue. | Charge distribution density. |
| GRAVY | Grand Average of Hydropathy (Kyte-Doolittle). | Hydrophobicity/solubility index Kyte and Doolittle (1982). |
| Aromaticity | Fraction of aromatic residues (F,Y,W). | Aromatic content affecting packing. |
| Instability_Proxy | Fraction of PEST/QD residues. | Proxy for protein stability Broom et al. (2017). |
| Hydro_Corr_L1 | Hydrophobicity autocorrelation lag 1. | Alternating hydrophobicity patterns. |
| Hydro_Corr_L2 | Hydrophobicity autocorrelation lag 2. | Periodic hydrophobicity ($\beta$-strands). |
| Hydro_Corr_L3 | Hydrophobicity autocorrelation lag 3. | Periodic patterns. |
| Hydro_Corr_L4 | Hydrophobicity autocorrelation lag 4. | $\alpha$-helix periodicity. |
| Hydro_Corr_L5 | Hydrophobicity autocorrelation lag 5. | Extended patterns. |
| Hydro_Corr_L6 | Hydrophobicity autocorrelation lag 6. | Long-range patterns. |
| Charge_Corr_L1 | Charge autocorrelation lag 1. | Alternating charge patterns. |
| Charge_Corr_L2 | Charge autocorrelation lag 2. | Periodic charge distribution. |
| Charge_Corr_L3 | Charge autocorrelation lag 3. | Periodic patterns. |
| Charge_Corr_L4 | Charge autocorrelation lag 4. | Helical charge periodicity. |
| Charge_Corr_L5 | Charge autocorrelation lag 5. | Extended patterns. |
| Charge_Corr_L6 | Charge autocorrelation lag 6. | Long-range patterns. |
| Shannon_Entropy | Sequence complexity (amino acid distribution). | Evolutionary conservation/variability Dill et al. (2007). |

#### 0.5 5. Constraint Frustration Features (11 features)

*Description: Per-residue conflicting interaction energies derived from variance in constraint graph weights. These features capture biophysical conflicts that serve as structural compatibility filters for challenging compartments.*

Table S5: Constraint Frustration Features (11 total)

| Feature Name | Description | Biological Interpretation |
|---|---|---|
| Frustration_NTerminal_Mean | Mean frustration in N-terminal region (residues 1-30). | Signal peptide vs. structural constraints. |
| Frustration_CTerminal_Mean | Mean frustration in C-terminal region (last 10 residues). | Retention signal vs. structural constraints. |
| Frustration_Structural_Mean | Mean frustration in middle structural region. | Core folding constraints. |
| Frustration_SignalVsStructure | Difference between N-terminal and structural frustration. | Signal region specificity indicator. |
| Frustration_MaxInSignalRegion | Maximum frustration value in N-terminal region. | Local constraint conflict hotspots. |
| Frustration_Localization | Inverse entropy of frustration distribution. | How localized vs. distributed conflicts are. |
| Frustration_HotspotCount | Number of residues with frustration ¿ mean + $2\sigma$. | Count of severe constraint conflicts. |
| Frustration_SatisfactionRatio | Proportion of residues with frustration ¡ median. | Fraction of residues with compatible constraints. |
| Frustration_SignalCorrelation | Correlation between frustration and sequence Signal scores. | Conservation vs. constraint conflicts alignment. |
| Frustration_HighSignal Frustration | Binary flag for high frustration in signal region. | Indicates significant signal-region conflicts. |
| Frustration_PerResidue_Mean | Mean of per-residue frustration vector (first 100 residues). | Summary of overall constraint conflicts. |

### Supplementary Material S2: Feature Summary and SHAP Framework

Table S6 provides a summary of the five feature categories. The SHAP (SHapley Additive exPlanations) framework Lundberg and Lee (2017) was used to explain model predictions, providing theoretically sound quantification of each feature's contribution.

Table S6: Summary of BioGraphX Feature Categories (Total: 157 features)

| Category | # Features | Scale | Biological Insight |
|---|---|---|---|
| Topological Graph | 85 | Residue to protein | Protein fold complexity, domain organization, network properties |
| Hybrid Interaction | 22 | Pairwise interactions | Specialized structural motifs, synergistic forces |
| Targeting Signal Features | 20 | Whole-protein | Known targeting signals + structural compatibility |
| Global Physicochemical | 19 | Whole-protein | Biophysical constraints, solubility, stability |
| Constraint Frustration | 11 | Per-residue | Conflicting interaction energies, structural compatibility filters |
| **Total** | **157** | **Multi-scale** | **Comprehensive sequence-structure-function representation** |

#### Supplementary Material S3: Computational Details

The time complexity for feature extraction is $O(n^2)$ for graph construction (where $n$ is sequence length), with optimizations for long sequences:

- **Short** ($\leq 2,000$ aa): Full processing
- **Medium** ($2,000 < L \leq 10,000$): Smart truncation (40% N, 20% middle, 40% C)
- **Long** ($> 10,000$ aa): Sliding window (window=1000, stride=500) with weighted aggregation

#### Supplementary Material S4: Parallel Implementation and Scalability

##### S4.1 Parallel Processing Architecture

The BioGraphX encoding pipeline employs a data-parallel architecture where independent protein sequences are processed concurrently. The implementation uses Python's Joblib library with the Loky backend for process-based parallelism. Key design decisions:

- **Embarrassingly parallel design**: Each protein sequence is encoded independently, enabling perfect scalability.
- **Dynamic batching**: Sequences are automatically grouped into batches sized according to available memory and core count.
- **Fault tolerance**: Worker processes are isolated to prevent single-sequence failures from affecting the entire batch.
- **Memory efficiency**: Large sequences (>10,000 aa) trigger adaptive processing before parallel distribution.
